# Epigenetic reprogramming towards mesenchymal-epithelial transition in ovarian cancer-associated mesenchymal stem cells drives metastasis

**DOI:** 10.1101/2020.02.25.964197

**Authors:** Huihui Fan, Huda Atiya, Yeh Wang, Thomas R Pisanic, Tza-Huei Wang, Ie-Ming Shih, Kelly K Foy, Leonard Frisbie, Chelsea Chandler, Hui Shen, Lan Coffman

**Affiliations:** Center for Epigenetics, Van Andel Research Institute, Grand Rapids, MI, USA; Division of Hematology/Oncology, Department of Medicine, Hillman Cancer Center, University of Pittsburgh, Pittsburgh, PA, USA; Department of Pathology, Department of Gynecology and Obstetrics, Department of Oncology, Johns Hopkins Medical Institutions, Baltimore, MD; Johns Hopkins Institute for NanoBiotechnology, Johns Hopkins University, Baltimore, MD, USA; Division of Gynecologic Oncology, Department of Obstetrics, Gynecology, and Reproductive Sciences, Magee Women’s Research Institute, University of Pittsburgh, Pittsburgh, PA

**Keywords:** Mesenchymal stem cells, carcinoma-associated mesenchymal stem cells, ovarian cancer, tumor microenvironment, epigenetic reprogramming, EZH2, WT1, metastasis, mesenchymal to epithelial transition

## Abstract

Ovarian cancer develops early intra-peritoneal metastasis establishing a supportive tumor microenvironment (TME) through reprogramming normal mesenchymal stem cells into carcinoma-associated mesenchymal stem cells (CA-MSCs). CA-MSCs are the stromal stem cell of the TME, supporting cancer growth, increasing desmoplasia, angiogenesis and chemotherapy resistance. We demonstrate epigenetic reprogramming drives CA-MSC formation via enhancer-enriched DNA hypermethylation, altered chromatin accessibility and differential histone modifications inducing a partial mesenchymal to epithelial transition (MET) increasing adhesion to tumor cells. Direct CA-MSC:tumor cell interactions, confirmed in patient ascites, facilitate ovarian cancer metastasis through co-migration. WT1, a developmental mediator of MET, and EZH2, mediate CA-MSC epigenetic reprogramming. *WT1* overexpression induces CA-MSC conversion while *WT1* knock-down, along with EZH2 inhibition, blocks CA-MSC formation. EZH2 inhibition subsequently decreases intra-abdominal metastasis.

**Significance:** This work presents a new paradigm of stromal reprogramming involving a partial mesenchymal to epithelial transition. Rather than a classic tumor cell epithelial to mesenchymal transition, metastasis relies on epigenetic rewiring of a CA-MSC allowing enhanced tumor cell binding and co-migration with tumor cells to form metastasis. Indeed, CA-MSCs in complex with tumor cells are abundant in patient ascites. Reversion of CA-MSCs to normal MSCs is observed in patients achieving complete response with neoadjuvant therapy. Identification of WT1 and EZH2 as mediators of the epigenetic reprogramming of CA-MSCs present potential targets to block the formation of CA-MSCs thus disrupting the TME and limiting ovarian cancer metastasis.

## Introduction

Ovarian cancer, the deadliest gynecologic cancer, kills over 14,000 US women yearly and 70% of all women diagnosed with the disease. This high mortality is due to early, diffuse intra-abdominal metastatic spread. Ovarian cancer quickly colonizes the peritoneal cavity building a complex tumor microenvironment (TME) that supports ovarian cancer survival, growth, and spread. This TME has unique physical and chemical properties with a complex bionetwork of tumor cells, immune cells, and stromal cells interacting within a hypoxic, nutrient poor environment (Zhang *et al*. 2003; Tothill *et al*. 2008; Verhaak *et al*. 2013; Karlan *et al*. 2014; Konecny *et al*. 2014).

Efforts to determine what drives the formation and function of the ovarian TME led to the identification of a critical stromal stem cell, the carcinoma-associated mesenchymal stem cell (CA-MSC) (McLean *et al*. 2011). CA-MSCs are benign stromal cells that meet all the criteria for mesenchymal stem cells/multipotent mesenchymal stromal cells (MSCs) as defined by the International Society for Cellular Therapy (ISCT) (Dominici *et al*. 2006). CA-MSCs are not tumor cells. Karyotype analysis, high resolution genotyping, and *in vivo* tumor initiation assays demonstrated that CA-MSCs have normal genomes, lack tumor-related mutations (such as p53 mutations), and isolated CA-MSCs lack malignant potential (McLean *et al*. 2011; Verardo *et al*. 2014). CA-MSCs are distinct from carcinoma associated fibroblasts (CAFs) and other more terminally differentiated stromal cells with long term proliferative capacity, lineage differentiation capacity and a unique gene expression pattern (Augsten 2014; Coffman *et al*. 2019). CA-MSCs strongly enhance ovarian cancer initiation, growth, chemotherapy resistance, increase the cancer stem cell-like (CSC) pool, and dramatically alter the stromal TME, increasing tumor-associated fibrosis and inducing angiogenesis (Spaeth *et al*. 2009; Cho *et al*. 2012; Coffman *et al*. 2016b). CA-MSCs have a unique gene expression profile characterized by altered extracellular matrix protein and cytokine expression (McLean *et al*. 2011; Coffman *et al*. 2016b; Coffman *et al*. 2019). CA-MSCs arise from normal tissue MSCs through cancer-mediated reprogramming (Coffman *et al*. 2019). Using a 6-gene mRNA expression classifier that accurately distinguishes normal MSCs from CA-MSCs, we demonstrated that ovarian cancer reprograms normal ovary and omental MSCs into ovarian cancer promoting CA-MSCs. Thus, ovarian cancer drives the formation of its own supportive TME (Coffman *et al*. 2019).

The mechanism driving this reprogramming remains unknown however given the wide-spread, durable expression changes without gain of genomic mutations we hypothesized epigenetic reprogramming likely drives this conversion. Cancer-related epigenetic changes are well documented, although most studies focus on epigenetic modification of cancer cells rather than associated stroma (Sharma *et al*. 2010; Poli *et al*. 2018).

Here we demonstrate that CA-MSCs have a unique epigenetic landscape compared to normal MSCs characterized by enhancer-enriched DNA hypermethylation, enrichment in repressive histone modifications and altered chromatin accessibility. We demonstrate DNA methylation changes are acquired during cancer-stimulated conversion of normal MSCs into CA-MSCs and are not altered by lineage differentiation into fibroblasts or adipocytes. However, MSCs derived from patients achieving a complete pathologic response to neoadjuvant chemotherapy revert to a normal MSC profile. Unexpectedly, the CA-MSC DNA methylation pattern partially resembles fallopian tube epithelium (FTE) while normal MSCs align more closely with fibroblasts. CA-MSCs appear to undergo a partial mesenchymal to epithelial transition with increase in cell contact pathways and functional demonstration of increased tumor cell adhesion. CA-MSCs increase ovarian cancer metastasis via direct tumor cell binding and co-migration in an orthotopic mouse model. CA-MSCs are also found in complex with tumor cells isolated from patient ascites. Further, we identify EZH2 and WT1 as potential mediators of CA-MSC epigenetic mesenchymal to epithelial reprograming. Interrupting this reprogramming dramatically decreases the ability of ovarian cancer cells to form intra-abdominal metastasis.

## Results

### CA-MSCs have a unique DNA methylation profile compared to normal tissue MSCs

We hypothesized that CA-MSCs undergo epigenetic alterations which drives their mitotically stable pro-tumorigenic phenotype. We used the Infinium MethylationEPIC (850K/EPIC/HM850) array to profile 13 CA-MSCs derived from high-grade serous ovarian cancer (HGSC), 20 omental/adipose/fallopian tube (FT) derived normal MSCs, and six patient tumor cell samples (sup table 1). We first performed QC on these samples, for both data quality (methods) and sample quality including purity determination and potential samples swaps. The promoter of MIR200C/141 is unmethylated in epithelial cells and methylated in mesenchymal cells (Vrba *et al*. 2010), and the methylation level (beta value) of MIR200C/141 directly corresponds to the fraction of non-epithelial cells (Cherniack *et al*. 2017; George *et al*. 2018). As expected, our MSCs (CA-MSCs and normal MSCs) had close to 100% methylation at the MIR200C/141 promoter site, consistent with a non-epithelial origin and high sample purity. In contrast, primary human tumor cell samples demonstrated lack of methylation at this locus (Fig1A). One CA-MSC sample had low MIR200C/141 promoter methylation consistent with tumor cell contamination and this sample also clustered with sorted tumor cells across the most variably methylated sites in the genome (SupFig1A) and was removed from all further analysis.

**Figure 1.**
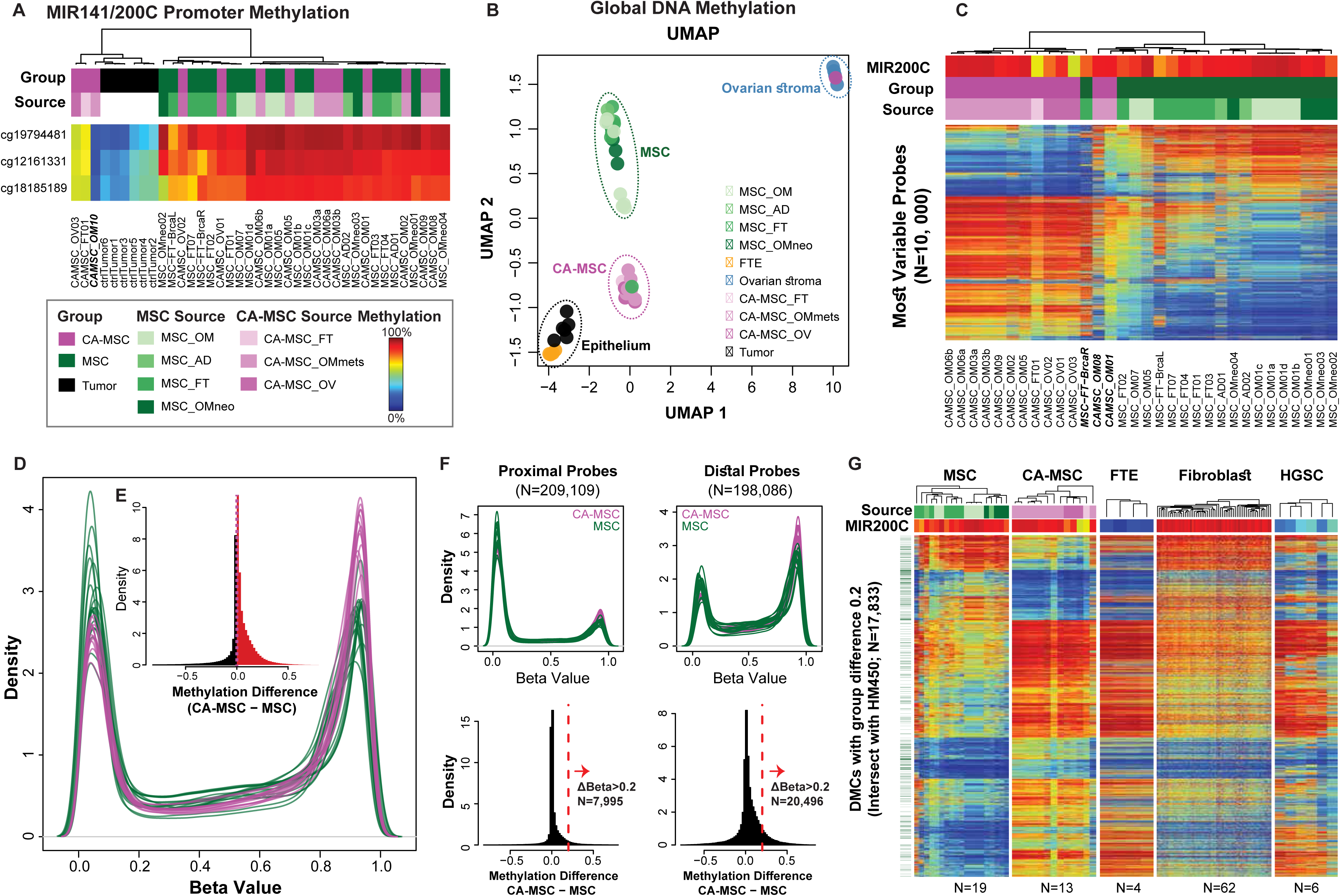
CA-MSCs have unique DNA methylation profiles compared to normal tissue MSCs. A. DNA methylation at the *MIR141/MIR200C* promoter separates samples of epithelial and mesenchymal nature. Each row of the heatmap shows a probe from the *MIR141/MIR200C* promoter region. Each column shows one of the CA-MSC, normal MSC (referred to as “MSC”) or tumor samples, with accompanying column-side color bars indicating the sample group (‘Group’), or isolation site (‘Source’, if MSC and CA-MSC) for each sample. A blue-to-red gradient indicates a beta value of 0-1 (DNA methylation level of 0% to 100%) in all heatmaps. B. Samples aggregate by cell type by DNA methylation. Normal fallopian tube epithelium and ovarian stroma samples are included as controls. Samples are shown as dots on scatterplots in the first two dimensions (x axis – Dimension 1; y axis – Dimension 2) from UMAP of the intersect set of 438,064 CpG loci present on both the EPIC and HM450 platform. Each cell type cluster is indicated by dotted circle with label. C. CA-MSCs have differential DNA methylation vs MSCs. Top variable CpG probes (N=10,000, filter with standard deviation) among 13 CA-MSC and 19 MSC samples, are shown as rows and samples are shown as columns, with annotations similar to Panel A. *MIR141/200C* promoter methylation (average of the three probes shown in A) for each sample is used to track the percentage of mesenchymal component proportion in each sample. D. Overall distribution (as smoothed probability density distribution) per sample of DNA methylation beta values (x axis) for CA-MSCs (purple lines) and MSCs (green lines). E. Histogram showing the distribution of DNA methylation (beta value) group differences between CA-MSCs and normal MSCs. Red represents sites with methylation difference > 0 (hypermethylation in CAMSCs), and black represents sites with methylation difference <0 (hypomethylation in CAMSCs). F. Distal regions show more frequent DNA methylation gain in CA-MSCs compared to MSCs. Both density and histogram plots are shown for proximal and distal CpG probe set (vertical in order). Proximal probes are defined as probes overlapping with transcript promoters (+/-500 bp around the TSS), while distal probes are defined as probes located near transcription factor binding sites (+/-100bp) but not within +/-2 kb away from the TSS. Distal elements are almost three times more likely to acquire a DNA methylation gain greater than 0.2 in CA-MSCs. G. DNA methylation heatmap showing differential methylation cytosine sites (DMC; N=17,883) between CA-MSCs and MSCs. Other tissue types are included for comparison. Visual similarities can be appreciated between MSCs and fibroblasts, and CA-MSCs to tissues of epithelial origin (ovarian tumor cells and fallopian tube epithelium/FTEs) respectively.

We downloaded and reprocessed external DNA methylation data on normal cell types from prior publications (Patch *et al*. 2015; Klinkebiel *et al*. 2016) (methods), including normal FTE and ovary stroma, profiled on the earlier generation of the Infinium platform, Infinium HumanMethylation450 (450K) array. The intersect between the two platforms (450,161 CpGs) were used in joint analysis with these external normal controls. Uniform Manifold Approximation and Projection (UMAP) reveals clustering coinciding with sample type (Fig1B), with a few exceptions: 1) One CA-MSC (CAMSC-OM08) grouped with the normal ovary stroma. After final pathology review, this sample was found to be derived from an ovarian carcinosarcoma, not HGSC. 2) One additional sample, CAMSC-OM01, grouped with CA-MSCs on the UMAP but clustered with the normal MSCs on the unsupervised hierarchical clustering (Fig1C) and overall demonstrated an intermediate profile between normal MSCs and CA-MSCs. This sample was derived from tumor adjacent omentum without clear malignant involvement and may represent a mixture of normal MSCs and CA-MSCs. 3) MSC-FT-BrcaR, derived from the right FT of a patient with a germline BRCA2 mutation undergoing risk reduction surgery grouped with CA-MSCs. In contrast, MSCs derived from the left FT of the same patient grouped with normal MSCs as expected. The expression-based CA-MSC classifier score for the left FT MSCs was 0.01 (consistent with a normal MSC) and for the right FT MSC was 0.95 (approaching the CA-MSC threshold of 0.97) in line with the methylation grouping and indicating certain stromal cells derived from a high risk patient more closely align with cancer associated stroma (Coffman *et al*. 2019). As expected, flow-sorted tumor cells were the closest to normal FTE, the presumed cell of origin for HGSC.

### MSCs derived from patients with complete response to chemotherapy resemble normal MSCs

Interestingly, MSCs derived from the omentum of patients treated with neoadjuvant chemotherapy who achieved a pathologic complete response clustered with the normal MSCs (n=4, MSC-OMneo1-4). Likewise, the expression classifier scores of these MSCs ranged from 0.01-0.15 (all within the normal MSC range). Radiographic evidence of omental involvement prior to neoadjuvant chemotherapy was documented in all cases.

### CA-MSCs demonstrate epigenetic alteration characterized by hypermethylation at distal enhancers

Top 10,000 most variably methylated probes across all MSC samples (CA-MSCs and normal MSCs) demonstrated a clear delineation between CA-MSCs and normal MSCs (Fig1C). CA-MSCs demonstrated distinct gains in DNA methylation vs normal MSCs (Fig1D, 1E). Greater than 128,000 differentially methylated cytosines (DMCs; 32% out of ∼401,000 filtered probes; methods), mapping to 22,991 differentially methylated regions (DMRs), were identified based on a threshold of FDR < 0.05. The most significant DMRs (ranked P-value followed by mean beta value levels) are listed in supplemental table 2. Such difference is more pronounced at distal rather than proximal sites (Fig1F; SupFig S2A,B), with distal regions three times more likely to acquire hypermethylation of >0.2 beta value difference compared to proximal regions consistent with DNA methylation changes preferentially occurring in enhancer regions.

### CA-MSC DNA methylation profile demonstrates similarity to that of epithelium

We examined sites with absolute beta value difference of greater than 0.2 in CA-MSCs vs normal MSCs (n=36,606 in HM850 loci, n=17,904 in HM850/HM450 intersect). Unexpectedly, in this space, CA-MSCs closely resembled FTE (Fig1G), which are benign epithelial cells, while normal MSCs were more similar to fibroblasts (of dermal origin). Similarly, CA-MSCs bore more resemblance to sorted HGSC cells, which are also epithelial in origin.

Intrigued by this result, we further examined all sites (instead of just CA-MSC vs MSC differential sites) in the HM850/HM450 intersect. Globally CA-MSCs exhibit a similarity to FTE (SupFig S1B), but are clearly not identical. We continued to investigate sites shared between, or unique to MSCs, CA-MSCs and FTE (methods; Fig2A). Normal ovarian stroma, fibroblast, high-grade serous ovarian cancer (HGSC), Sarcoma (SARC) and Uterine carcinosarcoma (UCS) are plotted as representatives of various relevant cell types. This new analysis revealed that CA-MSCs had a DNA methylation profile that combined MSC and FTE features, with a limited set of CA-MSC-specific hyper- and hypo-methylated sites (Fig2A). Thus, CA-MSCs demonstrated a partial conversion from normal MSC towards FTE. CA-MSCs are also distinct from other epithelial and mesenchymal malignancies including uterine carcinosarcoma and soft tissue sarcomas.

### Quantitative methylation specific PCR (qMSP) validates DMRs

To confirm the methylation differences noted in the EPIC array, and to develop an economical assay for classification, we selected a panel of five of the top DMRs (CA-MSCs vs MSCs): two regions of hypermethylation (*SASH1*, *C13orf45*) and three regions of hypomethylation (*ELN*, *GATA6*, *MARVELD2*). Average beta values for each CpG as determined by the microarray are plotted in Figure 2B. Quantitative MSP (qMSP) assays were designed for each locus. qMSP was performed on an independent set of 6 additional normal MSCs and 9 CA-MSCs. Across the five loci tested, the pattern of methylation between CA-MSCs and normal MSCs was confirmed with significant methylation demonstrated in CA-MSCs at *SASH1* and *C13Orf45* and significant hypomethylation at ELN, GATA6 and MARVELD2 (Fig2C).

**Figure 2.**
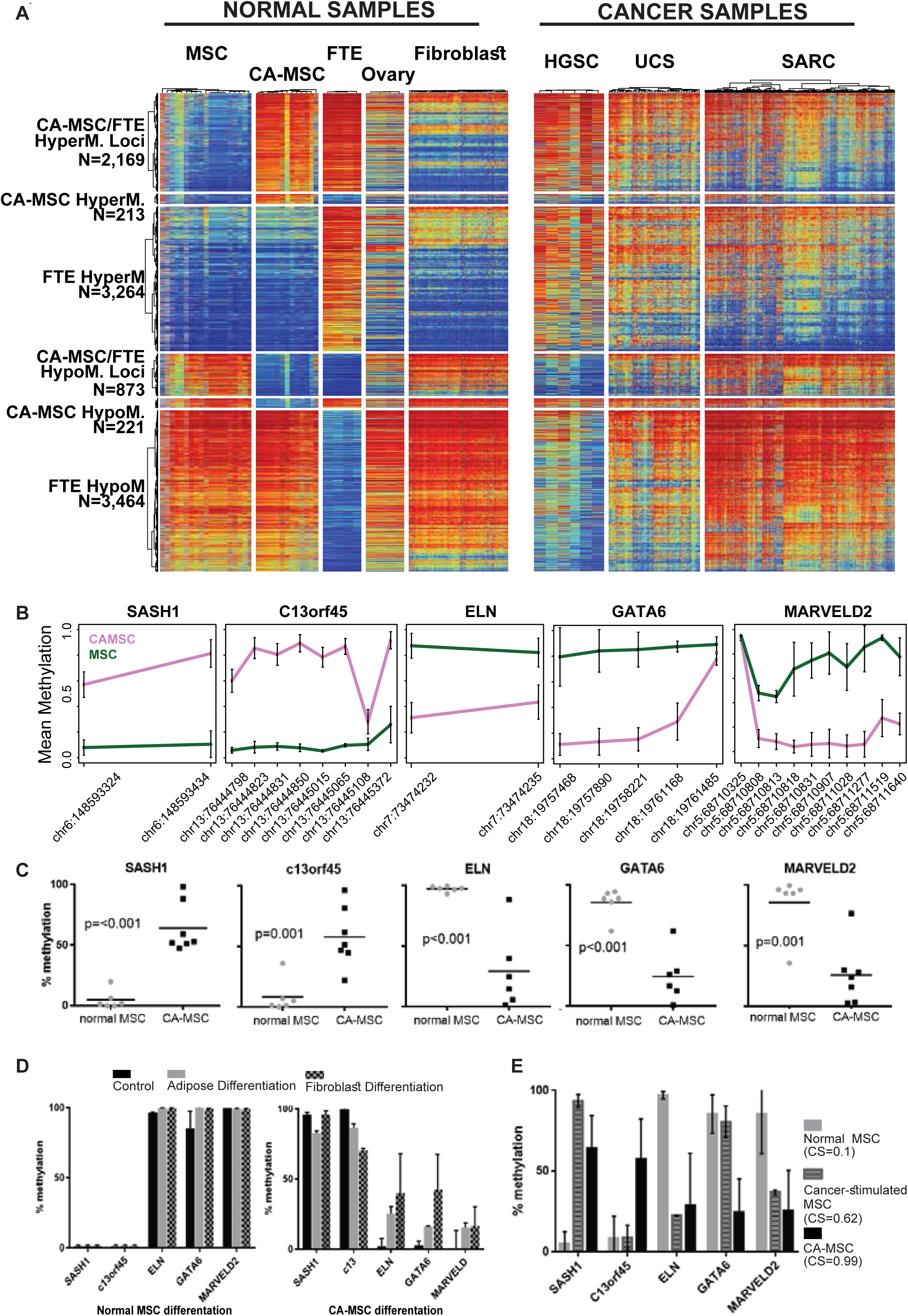
Verification of differential DNA methylation loci demonstrating methylation differences persist with MSC differentiation and are acquired during CA-MSC reprogramming. A. DNA methylation heatmap showing six different categories of CpG probes hyper- or hypo-methylated in different combinations of CA-MSC, FTE and MSC. Probe categories and the number of probes included are labelled on the left side of the heatmap. A blue-to-red gradient indicates a beta value of 0-1 (DNA methylation level of 0% to 100%) in all heatmaps. Corresponding cell types or cancer types are labelled on top of each heatmap panel. B. Top DMRs within five different genes chosen for quantitative methylation specific PCR (qMSP) validation. Mean beta values for CA-MSCs and MSCs are plotted on the y-axis against probe locations as plotted on the x-axis. Within-group variability is indicated by standard error bars for each group at each probe locus. C. qMSP validation of differential methylation between groups of CA-MSCs and normal MSCs shown in B, using independent CA-MSC and MSC samples. D. Both CA-MSCs and MSCs that undergo lineage differentiation (adipose and fibroblast differentiation) do not demonstrate significant DNA methylation changes at the selected loci as shown in C. Mean & SEM of 3 independent samples are represented. E. Cancer stimulation of normal MSCs *in vitro* induces a partial CA-MSC methylation pattern. MSCs that underwent partial conversion to a CA-MSC phenotype via hypoxic co-culture with ovarian cancer cells acquire CA-MSC-related methylation changes in three (*SASH1*, *ELN*, *MARVELD2*) out of the five loci tested. Mean & SEM of 3 independent samples are represented.

### CA-MSC-associated DMRs persist throughout lineage differentiation and are acquired during cancer-mediated reprogramming of MSCs

To determine if lineage differentiation alters methylation at the identified DMRs, we also used qMSP to characterize normal MSC and CA-MSC *in vitro* lineage directed differentiation into fibroblasts or adipocytes (Fig2D). Interestingly, these cells maintained their methylation status at these loci even after differentiation.

Given our previous work demonstrating that normal MSCs are induced by tumor cells to become CA-MSCs, we analyzed *in vitro* indirect cancer stimulation of normal MSCs (which yields a partial conversion to a CA-MSC) for the development of CA-MSC specific DMRs with the qMSP assays developed above. Three of the five loci tested demonstrated CA-MSC-like alterations, with acquisition of DNA methylation at *SASH1* and loss of methylation at *ELN* and *MARVELD2*. *C13orf45* and *GATA6* loci remained unchanged (Fig2E). This finding suggests a partial conversion to a CA-MSC DNA methylation profile with *in vitro* cancer stimulation. This is consistent with a change in the CA-MSC classifier score from 0.1 in normal MSCs (consistent with a normal phenotype) to 0.62 after indirect cancer stimulation (threshold for CA-MSC is 0.97).

### CA-MSCs have altered chromatin accessibility

We next analyzed the chromatin accessibility of CA-MSCs vs normal MSCs using the Assay for Transposase-Accessible Chromatin with high-throughput sequencing (ATACseq). As expected, clustering based on ATACseq peaks accurately separated normal MSCs from CA-MSCs, with two replicates from the same sample located right next to each other as expected for such controls (Fig3A). DESeq2 identified 5,129 statistically differential peaks with an adjusted p value lower than 0.05. Power was limited in this analysis, partly due to the small sample size after removing samples that failed QC. However, even in the majority of sites that did not reach statistical significance, CA-MSCs in general demonstrated a decrease in peak strength compared to MSCs after normalizing for library size (Fig3B), consistent with the observed widespread gain of DNA methylation in CAMSCs. The majority of the differential peaks between CA-MSCs and MSCs were found in distal intergenic regions, closely tracking the DNA methylation results (SupFig S2C,D). Overall, sites with reduced peak strength in CA-MSCs tended to have lower gene expression (Fig3C; R^2^=0.02; p value < 2.2e-16) and higher DNA methylation (Fig3D).

**Figure 3.**
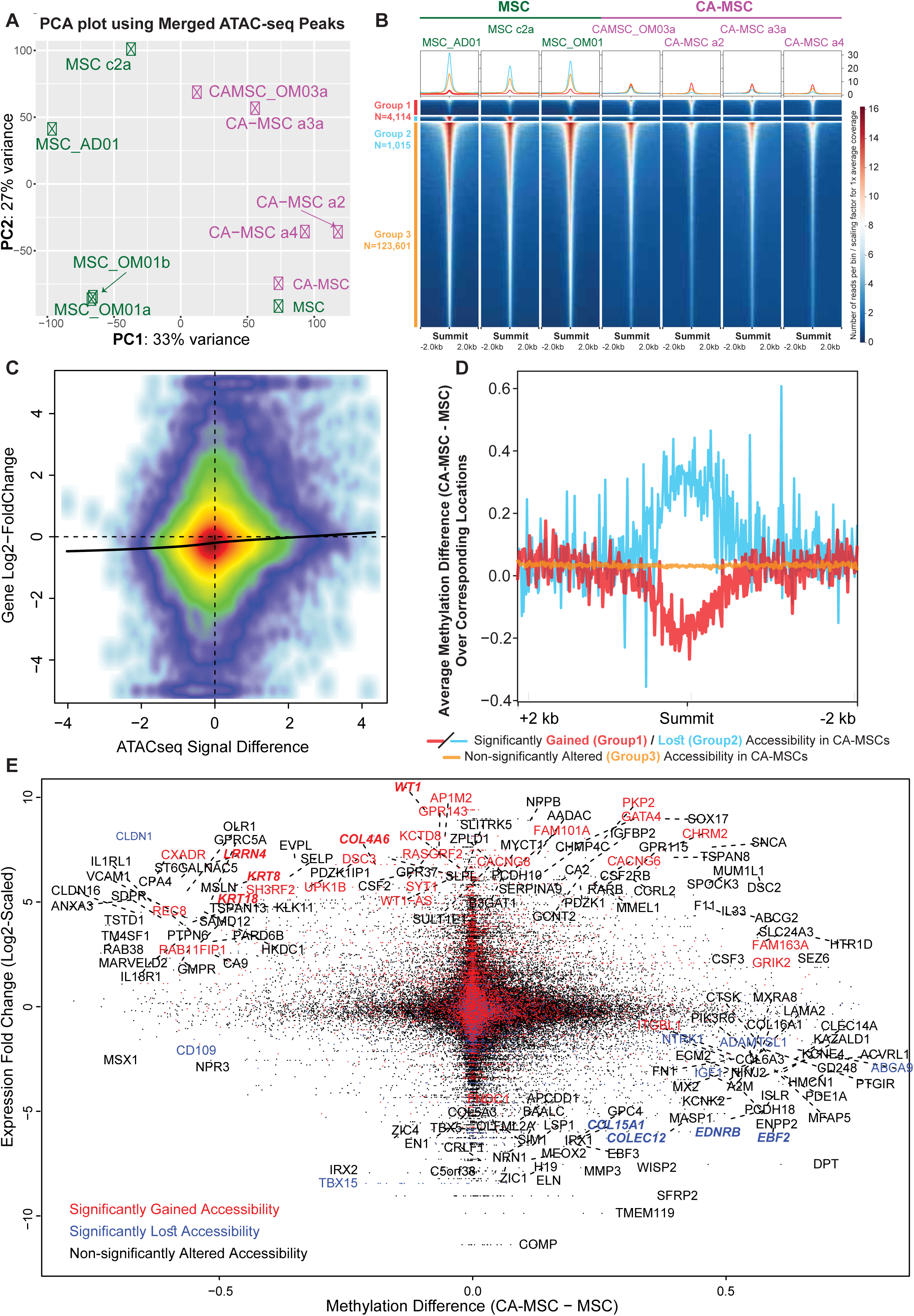
CA-MSCs have altered chromatin accessibility compared to normal MSCs which correspond to DNA methylation and RNA expression changes. A. Principal component analysis (PCA) on the merged peak regions from Assay for Transposase Accessible Chromatin with high-throughput sequencing (ATAC-seq) separates CA-MSCs from MSCs. B. Heatscatter plot showing the correlations between ATAC-seq peak signal difference (x axis) and their corresponding gene expression fold changes (y axis, log2-transformed), with color indicating local smoothed density. A LOWESS (Locally Weighted Scatterplot Smoothing) regression line is shown in black. C. ATAC-seq heatmaps centered at the summits of all the merged peaks, grouped into three categories: peaks (statistically) significantly gained (G1), lost (G2), and not with statistical significance (G3) in CA-MSCs comparing to MSCs. Heatmap is colored based on 10 bp-binned ATAC-seq signal, normalized against sequencing library size per sample. D. DNA methylation patterns around the summits for peak group G1, G2, and G3 shown in C. Line plot is centered on the summits of all the peaks within each group plotted as x-axis, while average of the overlapping methylation difference is shown as y-axis. E. Integration of DNA methylation, gene expression, and ATAC-seq calls. Probe-level DNA methylation difference (CA-MSC - MSC) for promoter probes (defined as +/-1.5 kb away from TSS) is plotted as x-axis. Expression fold changes (log2-scaled) for the corresponding genes are plotted as y-axis. Dots are colored based on gene-overlapping ATAC-seq peak orientations, where red represents open, blue closed, and black unaltered peaks in comparing CA-MSCs to MSCs. Gene labels are colored the same way. Top candidate genes with largest gene expression alterations (50 each way based on the gene fold change rank), and combined methylation and gene expression differences (absolute beta value difference > 0.5 and fold change > 2) are labelled with gene names. Exemplified up and down genes are highlighted as bold and italic.

### DNA methylation and chromatin accessibility correspond to transcriptomic differences in CA-MSCs vs MSCs

We incorporated previously reported RNA sequencing data (Coffman *et al*. 2019) with our DNA methylation and chromatin accessibility data (Fig3E). In general, genes with gained peaks and genes with lost peaks segregated based on gene expression. Genes exemplified by *WT1*, *LRRN4*, *KRT8*, *KRT18* and *COL4A6* were significantly overexpressed in CA-MSCs and demonstrated hypomethylation and open ATAC peaks. In contrast, genes such as *COL15A1*, *EBF2*, *EDNRB*, and *COLEC12* which have decreased expression in CA-MSCs demonstrated hypermethylation and reduced ATAC peaks.

### Gene expression pathway analysis confirms CA-MSCs undergo a partial mesenchymal to epithelial transition (MET)

Pathway analysis of the most differentially expressed genes demonstrated an upregulation of multiple pathways associated with cell:cell and cell:matrix adhesion, a largely epithelial feature, and down-regulation of genes maintaining mesenchymal cellular identity (Fig4A-B). Genes downstream of the TGF-beta signaling pathway, a strong inducer for EMT (Xu *et al*. 2009), were significantly enriched in the downregulated set in CA-MSCs. We took top 25 epithelial (E-) and mesenchymal (M-) signature genes from a previous study (Creighton *et al*. 2013) ranked based on the Pearson’s correlations of gene expression with the EMT score calculated from the canonical EMT markers (Lee *et al*. 2006). CA-MSCs clearly exhibited higher expression for the E-genes and lower expression for the M-genes. A quantitative EMT score derived with the 16-gene signature from the same study also suggested that CA-MSCs indeed had a more “epithelial” signature while normal MSCs had a more “mesenchymal” signature (Fig4C). This observation coalesces with the DNA methylation results, which also suggest a partial mesenchymal-epithelial transition in CA-MSCs.

**Figure 4.**
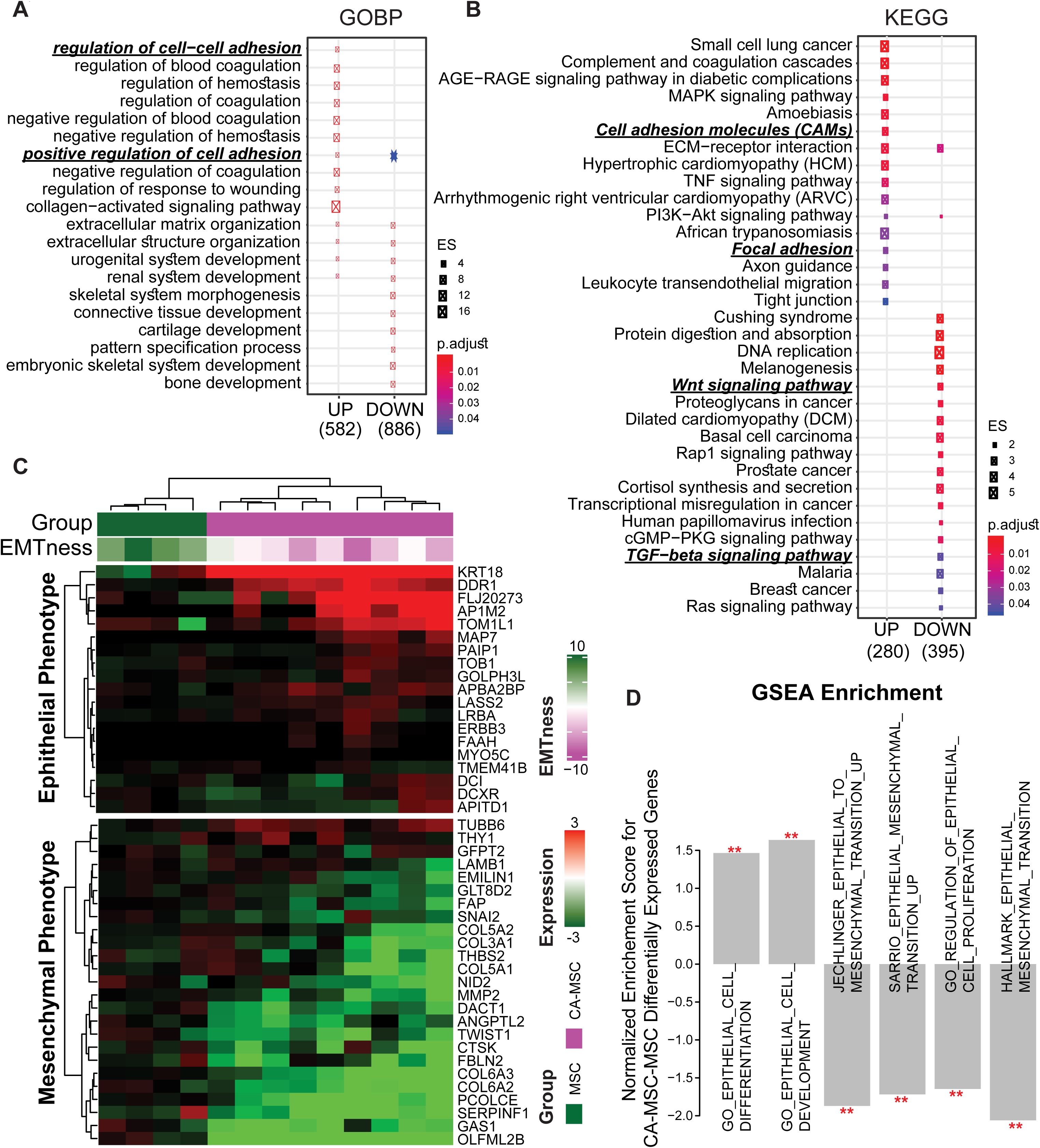
CA-MSCs undergo a partial mesenchymal to epithelial transition (MET). A. Gene ontology (GO) enrichment analysis (biological concept of Biological Process; GOBP). The numbers of up- and down-regulated genes comparing CA-MSCs to MSCs are labeled underneath the plot. Each dot represents a GOBP term, with dot size indicating enrichment score (ES), and dot color representing significance level. Two cell adhesion related processes are highlighted for genes upregulated in CA-MSCs. B. KEGG (Kyoto Encyclopedia of Genes and Genomes) pathway analysis. Two pathways relevant to cell adhesion are highlighted for upregulated genes, and two pathways relevant to EMT (epithelial-mesenchymal transition), the reverse process of MET, are highlighted for downregulated genes. C. Heatmap of epithelial- and mesenchymal-signature genes indicates a partial MET in CA-MSCs. EMTness score based on a 16-gene scoring system (higher and positive score indicate a more mesenchymal phenotype) is shown as column annotation, together with sample group information. D. Gene Set Enrichment Analysis (GSEA) results are in line with a potential MET in CA-MSCs. Genes are ranked based on fold changes comparing CA-MSCs to MSCs. Relevant GSEA terms are plotted as bars, with significance levels indicated by red stars. Normalized enrichment scores are plotted as y-axis.

To test if CA-MSCs exhibit an increase in epithelial phenotypic characteristics such as increased cell adhesion capacity compared to normal MSCs, we performed *in vitro* cell adhesion assays of tumor cells to normal MSCs or CA-MSCs. CA-MSCs had a 3-fold increase in tumor cell binding capacity after 30 mins of co-incubation. This pattern is true regardless of MSC or CA-MSC origin (fallopian tube or omentum) and is consistent across two ovarian cancer cell lines and one primary patient tumor cell line tested (Fig5A, SupFig3A). The increased tumor cell binding confirms the pathway analysis showing increased cell:cell adhesion in CA-MSCs.

**Figure 5.**
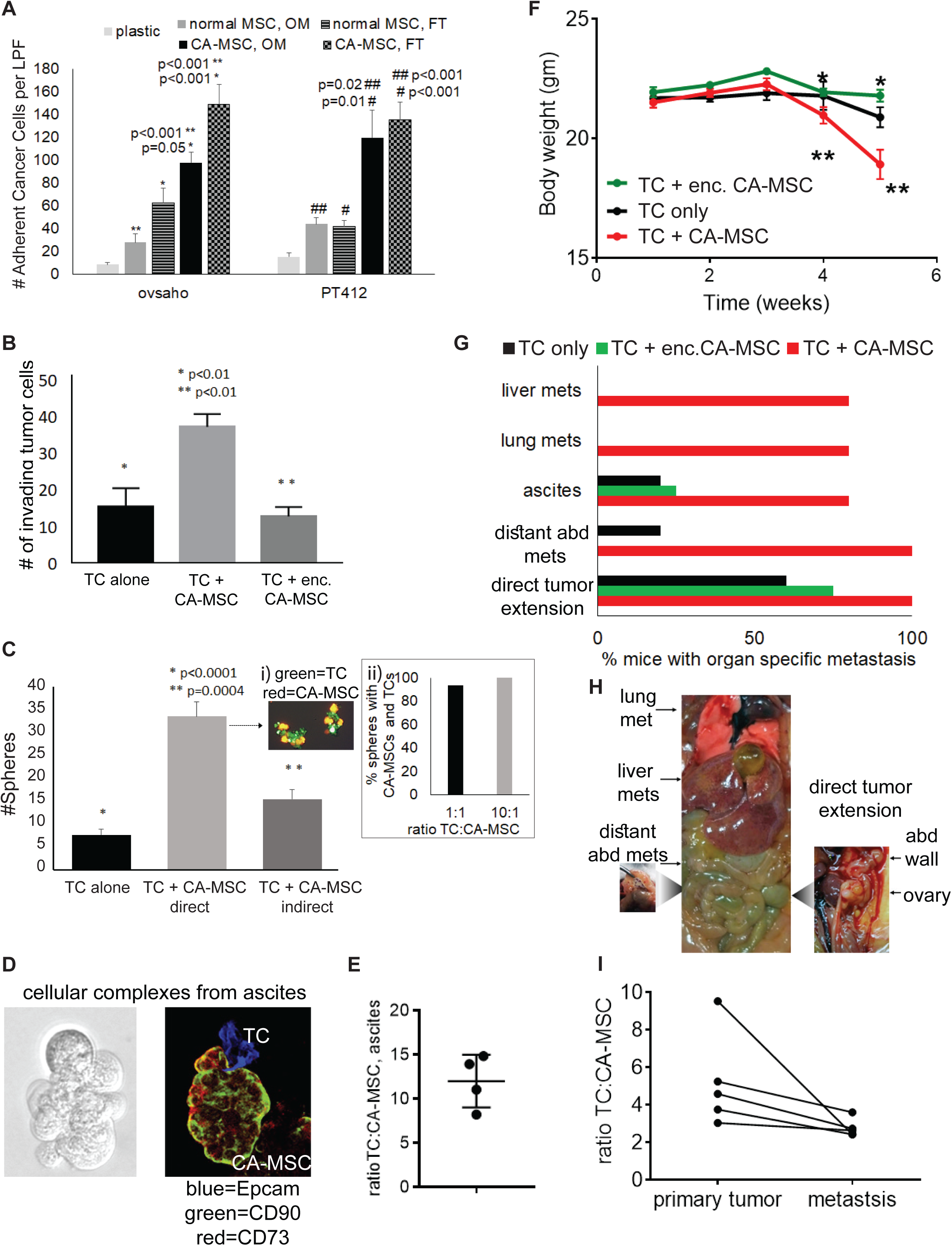
CA-MSCs have enhanced tumor cell binding and increase ovarian cancer metastasis through direct tumor cell:CA-MSC interactions and co-migration. A. CA-MSCs demonstrate greater cancer cell adhesion compared to normal MSCs. Quantification of the number of adherent cancer cells (OVSAHO or primary patient cancer cells pt412, GFP labeled) to plastic, normal MSCs derived from the omentum or fallopian tube (FT) or CA-MSCs derived from omental tumor or fallopian tube tumor via fluorescent microscopy, cell counts per low power field (LPF). OVCAR3 binding presented in supplemental Figure 3A. B. CA-MSCs enhance the invasion of OVSAHO tumor cells (TCs) through Matrigel coated transwells. Results were quantified via fluorescent microscopic counting of TCs which invaded to the underside of the transwell per LPF; TCs are GFP labeled, CA-MSCs are mT labeled. Alginate encapsulation (enc.) prevents enhanced invasion demonstrating the necessity of direct CA-MSC:TC contact. C. CA-MSCs enhance OVSAHO TC sphere growth under non-adherent condition through direct CA-MSC:TC interactions. GFP-TC and mT-CA-MSCs grown in low attachment plates were allowed to directly interact or were separated by a porous membrane. Spheres containing GFP-TCs were quantified via fluorescent microscopy per LPF and compared to TCs alone and TCs allowed to directly or indirectly interact with CA-MSCs. (i) representative picture of GFP-TCs and mT-CA-MSCs within heterocellular spheres. (ii) Quantification of the percent of spheres containing at least one CA-MSC per TC sphere at 1:1 and 10:1 TC:CA-MSC ratio demonstrating preferential formation of heterocellular spheres even at low CA-MSC concentrations. For all in vitro experiments, mean & SEM from 3 independent experiments are represented. D. Cellular complexes isolated from high grade serous ovarian cancer patient ascites. Confocal microscopy demonstrates TCs (as identified by Epcam+) in contact with CA-MSCs (identified with CD90+/CD73+/Epcam-). E. Quantification of the average ratio of TC and CA-MSCs within patient ascites cellular complexes. Cellular complexes were isolated from ascites, dissociated into single cells and TCs and CA-MSCs quantified by flow cytometry. Ratios from four independent patient ascites are presented. F-G. Orthotopic mouse model demonstrating increased metastasis with CA-MSCs in physical contact with TCs. Ovarian cancer cells were injected into the ovarian bursa alone (TC only), injected with CA-MSCs (TC + CA-MSC), or injected with encapsulated CA-MSCs (TC + enc. CA-MSC) and monitored for disease growth. F. Mouse weight over time across experimental groups demonstrates the TC + CA-MSC group had the most weight loss associated with systemic illness and disease burden (*, ** indicate significant decrease in weight between time points). G. Quantification of organs involved with gross metastatic disease at necropsy demonstrating increased lung, liver and intra-abdominal metastasis in the TC + CA-MSC group. H. Representative images of metastatic sites, including lung, liver, ovary, abdomen metastasis. I. CA-MSCs co-localize with TCs at the primary and metastatic tumor and metastatic tumor sites have an increased proportion of CA-MSCs. Quantification of the TC to CA-MSC ratio as measured via flow cytometry in the TC + CA-MSC group.

### CA-MSCs are found in direct contact with ovarian cancer cells in patient ascites

To understand the physiologic relevance of a partial MET resulting in enhanced adhesion of CA-MSCs to tumor cells, we isolated cellular complexes from ovarian cancer patient ascites. Flow cytometry analysis of complexes demonstrated the presence of CA-MSCs in a roughly 1:10 ratio to tumor cells (Fig5D). CA-MSC identity was verified via flow cytometry analysis with cell surface markers (CD90, 73, 105+/CD45, 34, 14, 19-) and differentiation capacity (adipocyte, osteocyte, chondrocyte) was confirmed to meet ISCT criteria for MSCs (Dominici *et al*. 2006). Cytospins of cellular complexes were analyzed with immunofluorescent confocal microscopy for Epcam (pacific blue) positive tumor cells and CD90 (FITC)/CD73 (RFP) positive CA-MSCs confirming physical interactions between CA-MSCs and tumor cell in ascites (Fig5D).

### CA-MSCs enhance ovarian cancer metastasis

As transcoelomic metastasis is thought to be a main mechanism of ovarian cancer metastasis resulting in diffuse peritoneal disease and malignant ascites coupled with the fact that CA-MSCs are found within cellular complexes in ascites, we next assessed the role of CA-MSCs in ovarian cancer metastasis. We tested if CA-MSCs support critical steps in ovarian cancer metastasis including migration/invasion and survival under non-adherent conditions. CA-MSCs allowed to directly interact with tumor cells (CAOV3, OVSAHO, OVCAR3) significantly enhanced tumor cell migration and invasion through a matrigel-coated transwell (Fig5B). CA-MSCs also enhanced the ability of tumor cells to grow under non-adherent conditions as spheres (Fig5C). Fluorescent microscopy of CA-MSCs (expressing mT) and tumor cells (expressing GFP) demonstrated CA-MSCs and tumor cells remained in physical contact during migration/invasion and sphere formation indicating co-migration and formation of heterocellular spheres (Fig5Ci). Increasing the ratio of tumor cells to CA-MSCs from 1:1 to 10:1 under non-adherent conditions resulted in each tumor cell sphere containing at least one CA-MSC even at low CA-MSC concentrations (Fig5Cii). This indicates a preferential direct binding between these cell types. This also mirrors the ratio of tumor cells to CA-MSCs (∼10:1) found in patient ascites cellular complexes (Fig5E).

### Direct tumor cell:CA-MSC interactions mediate enhancement of ovarian cancer metastasis

To confirm the enhancement of metastatic steps is mediated through direct CA-MSC:TC interactions, we repeated the above experiments but prevented the direct interaction between CA-MSCs and tumor cells by encapsulating CA-MSCs in alginate or separating the cells by a permeable membrane (allowing paracrine signaling without direct contact). Luciferase-secreting CA-MSCs were used to verify the continued viability and effective protein secretion of encapsulated CA-MSCs (SupFig3B). Alginate encapsulated CA-MSCs failed to enhance tumor cell migration/invasion (Fig5B). Similarly, CA-MSCs separated from tumor cells by a permeable membrane did not enhance tumor cell sphere formation (Fig5C). Together, this implicates direct contact between CA-MSCs and TCs as critical to *in vitro* enhancement of metastasis.

We next assessed the impact of CA-MSCs in a murine model of metastasis. Ovarian cancer cells (OVCAR3) were implanted orthotopically in the ovarian bursa alone, with CA-MSCs or with alginate encapsulated CA-MSCs. Mice with tumor cells + CA-MSCs demonstrated significant increases in metastasis and systemic illness as measured by decreased weight over time compared to mice with tumor cells alone or encapsulated CA-MSCs (Fig5F). Mice with tumor cells allowed to directly contact CA-MSCs demonstrated significantly more intra-abdominal metastasis with co-localization of tumor cells and CA-MSCs at metastatic implants consistent with tumor cell:CA-MSC co-migration (Fig5I). Mice with tumor cells alone or with encapsulated CA-MSCs demonstrated large localized ovarian tumors but did not develop ascites, distant abdominal metastasis, lung or parenchymal liver metastasis (Fig5H). CA-MSCs within the alginate capsule remained viable and were re-isolated at the time of necropsy. Collectively, this indicates CA-MSCs significantly enhance ovarian cancer metastasis and the epigenetically driven partial MET allowing increased tumor cell adhesion is critical to the pro-metastatic phenotype of CA-MSCs.

### *WT1* is a driver of MET and mediates CA-MSC formation

One of the most significantly differentially regulated genes with increased RNA expression, decreased DNA methylation and increased ATAC peak was *WT1* (Fig3E; Fig6A). *WT1* expression is often found in cells going through mesenchymal to epithelial transitions or cells with mixed epithelial and mesenchymal properties, such as podocytes (Moore *et al*. 1998). In addition, *WT1* overexpression has been reported to induce MET and is a regulator of MET during development (Hohenstein and Hastie 2006; Essafi *et al*. 2011). While known as a tumor marker, WT1 is present in tumor-associated stroma (Hylander *et al*. 2006; He *et al*. 2008). Independent qRT-PCR validation of *WT1* expression demonstrated ∼100 fold increased expression in CA-MSCs vs normal MSCs with corresponding increase in protein level (though protein levels varied more substantially than RNA expression) (Fig6B,C). Further, during cancer stimulated reprogramming of MSCs into CA-MSCs, *WT1* expression was induced (Fig6C).

Interestingly, while Figure 3E showed a decrease in *WT1* promoter methylation associated with WT1 overexpression, upon close examination of the entire *WT1* region (Fig6A), we discovered that the promoter DNA methylation differences were carried by only a subset of the normal MSC samples. However, two regions (highlighted by orange and cyan boxes in Figure 6A) in the gene body of *WT1* were consistently methylated in normal MSCs and unmethylated in CA-MSCs. Further, each of these differentially methylated regions overlapped with a site with differential accessibility between CA-MSCs and MSCs as determined by ATAC-seq. These ATAC-seq differential peaks were statistically significant even with the small sample size. In particular, when we looked up ENCODE histone modification data, Region 2 (cyan box) clearly functions as an enhancer in K562 cells, a malignant leukemia cell line, but is not active in normal fibroblasts (NHLF), a cell type normal MSCs appear to resemble based on DNA methylation results. These results seem to suggest that hypomethylation at this cryptic enhancer, associated with enhanced accessibility, likely turned this region normally methylated in normal MSCs into an active enhancer, and greatly promotes WT1 expression (>1,000 fold by RNA-seq, >100 fold by RT-PCR). Given the global trend of ***hyper***methylation in CA-MSCs, this specific ***hypo***methylation is even more likely to be a driver event, instead of a passenger event associated with global gain of DNA methylation.

**Figure 6.**
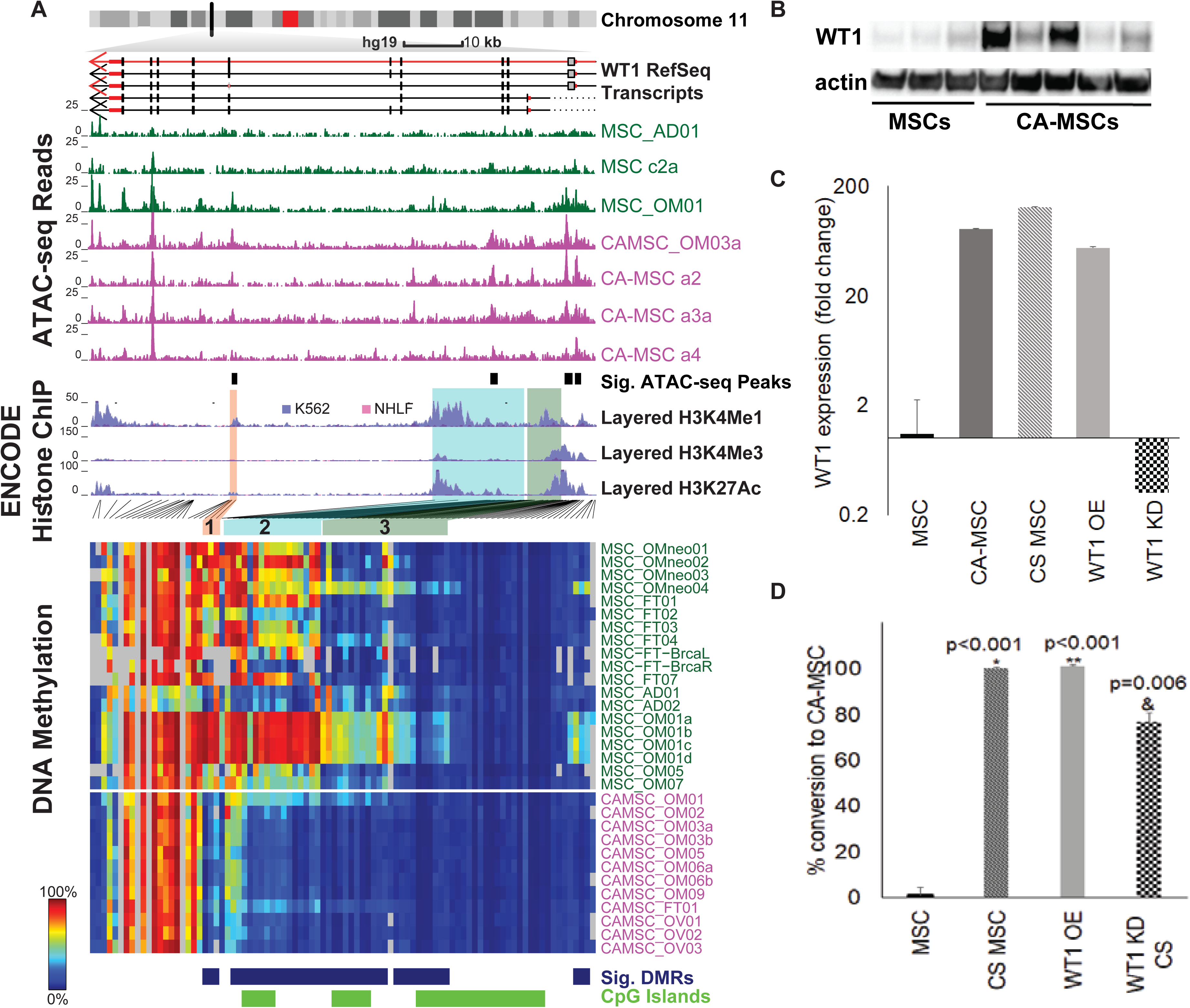
*WT1* is associated with a permissive epigenetic state in CA-MSCs and participates in the formation of a CA-MSC. A. Integrated view of epigenetic profiles for *WT1*. Normalized read count (10-bp bin) for ATAC-seq is plotted on top according to the genomic location they map, each row representing one sample. Significantly gained peaks in CA-MSCs are indicated by black boxes below the ATAC-seq signal plots. *WT1* transcripts are plotted under the chromosome ideogram, with arrows indicating the direction of the transcript. Layered H3K4me1 (enhancer mark), H3K4me3 (promoter mark), H3K27ac (active chromatin mark) ChIP-seq tracks from ENCODE for two cell types, K562 (a bone marrow derived myeologenic leukemia cell line, purple) and NHLF (normal human lung fibroblast, pink) are plotted below. WT1 transcripts, ATAC-seq peaks, and ENCODE histone ChIP-seq tracks are drawn to proportion on the hg19 genome with scale indicated on this panel. DNA methylation level for each CpG (columns of heatmap) in this region is plotted below as a heatmap. As the DNA methylation probes are not evenly distributed, this heatmap is not directly aligned with genomic locations for tracks above, but with lines connecting each CpG to the corresponding genomic location. Each row represents a sample as labelled on the right. Differentially methylated regions (DMRs) called by DMRcate between MSCs and CA-MSCs are indicated below with blue boxes (blue = DMR probe), followed by CpG island (CGI) designation for each probe (green=CGI probe). Three regions of interest discussed in the main text are highlighted with orange (Region 1), cyan (Region 2) and green (Region 3) shaded boxes. The locations for these regions are indicated again for the heatmap below with the same color. Region 1 and Region 2 are consistently methylated in normal MSCs and unmethylated in CAMSCs, while Region 3 overlapping the promoter region is only methylated in a subset of MSCs although a statistically significant DMR. B. Western blot of WT1 from MSCs (3 independent samples) and CA-MSCs (5 independent samples) demonstrating higher protein levels of WT1 in CA-MSCs, Actin is the loading control. C. Expression fold change of *WT1* vs normal fallopian tube MSC, CA-MSCs, CS MSCs, *WT1* overexpression (OE) MSCs and *WT1* siRNA knock down (KD) MSCs. D. Quantification of conversion to a CA-MSC in CS siRNA scrambled control MSCs, *WT1* OE MSCs alone, and CS *WT1* KD MSCs. CS was performed with hypoxic direct cancer cell co-culture. Values are normalized to the conversion of scrambled control MSCs based on the CA-MSC classifier score. Mean & SEM of 3 independent experiments is represented. *,** represents marked group vs. scrambled control without CS; & represents marked group vs. scrambled control with CS

To determine if WT1 is important in the formation of CA-MSCs, we created *WT1* overexpressing (OE) normal MSCs via lentiviral transduction. Compared to lenti-GFP control MSCs, *WT1* expression increased to levels approximating CA-MSCs (Fig6C). Analysis of the CA-MSC classifier score in *WT1* OE MSCs vs control MSCs demonstrated increase in the classifier score from 0.01 (a normal MSC score) to 0.99 (a CA-MSC score). This indicates WT1 OE drives the conversion of a normal MSC to a CA-MSC. We also performed siRNA mediated knock down (KD) of WT1 in normal MSCs. *WT1* KD MSCs underwent cancer stimulation via hypoxic direct co-culture with ovarian cancer cells which we previously demonstrated was the most effective *in vitro* condition to convert a normal MSC to a CA-MSC. Cancer-stimulated (CS) *WT1* KD, compared to CS scrambled control MSCs, demonstrated a 23% decrease in the conversion into a CA-MSC (based on change in CA-MSC classifier score) (Fig6D).

### CA-MSC expression pattern correlates with altered EZH2 binding sites

DNA methylation alterations at enhancer regions can serve as a readout of overall activity of transcription factors, where hypermethylation often indicates an inactive transcription factor binding site (TFBS), and hypomethylation likely marks an active state (Yao *et al*. 2015). Enrichment analysis for annotated TFBS sites identified EZH2 binding sites as the most significantly enriched category with hypermethylation in CA-MSCs (Fig7A). Concordantly, binding sites of SUZ12, a member of the polycomb repressive complex 2 (PRC2) along with EZH2, were also enriched.

**Figure 7.**
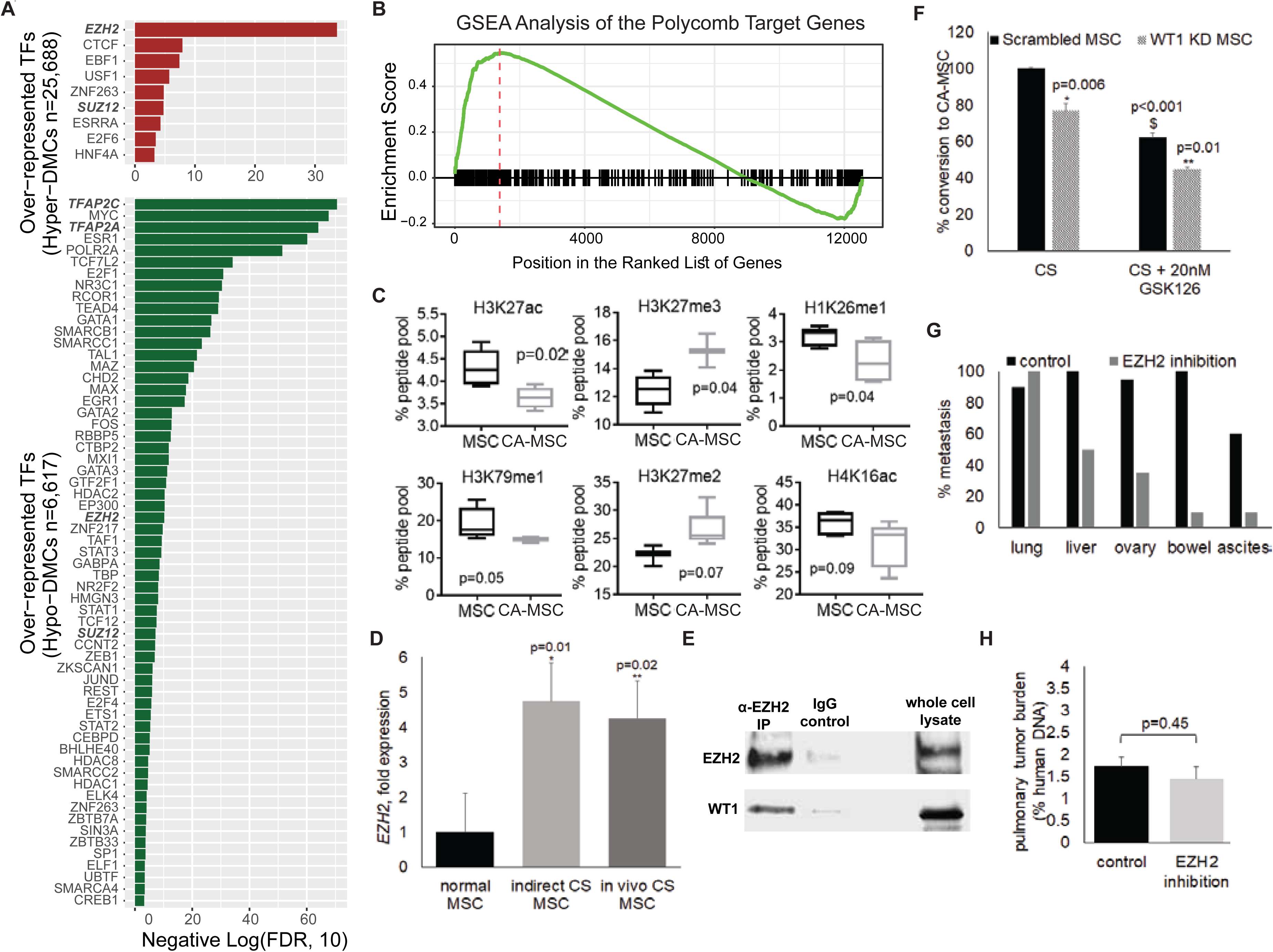
EZH2, along with WT1, impacts CA-MSC reprograming and ovarian cancer metastasis. A. Transcription factor binding sites (TFBS) enrichment for distal DMCs hypermethylated (top) and hypomethylated (bottom) in CA-MSCs. Negative log10-transformed FDRs are plotted as x-axis. Bolded gene names are referenced in text. B. GSEA plot shows enrichment of the Polycomb group (PcG) target genes on both ends of the ranked gene list (ordered by gene fold changes in comparing CA-MSCs to MSCs). C. Box plots of top histone modifications differentially represented in CA-MSCs vs. MSCs identified by mass spectrometry. D. *EZH2* expression is increased during *in vitro* (indirect) and *in vivo* cancer stimulation with ovarian cancer cells. Fold change compared to normal MSC values. Mean & SEM of 3 independent experiments is represented. E. EZH2 directly interacts with WT1. Co-immunoprecipitation of EZH2 pulls down WT1. F. *WT1* knock down (KD) in combination with EZH2 inhibition (EZH2i) partially blocks the formation of a CA-MSC. Quantification of the percent of conversion to CA-MSC in *WT1* KD MSCs or scrambled control after cancer stimulation (CS) with hypoxic direct cancer cell co-culture with, and without GSK126. Values are normalized to scrambled control conversion based on CA-MSC classifier score. Mean & SEM of 3 independent experiments is represented.*, ** represent comparison to scrambled control; & represents comparison to CS scrambled control without GSK126 G. EZH2 inhibition with GSK126 decreases the formation of intra-abdominal metastasis. The percent of mice developing metastasis at each organ site is represented, N=10 mice per group. H. EZH2 inhibition did not impact tumor cell growth in the lung as demonstrated by quantitative human vs. mouse specific PCR from extracted lungs.

Interestingly, EZH2/SUZ12 binding sites were also enriched in loci with hypomethylation (Fig7A), indicating a likely redistribution of EZH2/SUZ12. Consistent with this observation, gene set enrichment analysis (GSEA) demonstrated an enrichment in EZH2 targets in both up- and down-regulated genes in CA-MSCs compared to normal MSCs (Fig7B).

### CA-MSCs are enriched with PRC2-related histone marks

Given the role of EZH2 in histone modifications at H3K27, we investigated global changes in histone modifications between CA-MSCs and normal MSCs. We performed mass spectrometry on the most common 80 histone modifications in CA-MSCs and MSCs. Out of the 80 modifications tested, six histone marks were differentially represented in CA-MSCs vs MSCs (p value <0.1): H3K27ac, H3K27me3, H1K26me1, H3K79me1, H3K27me2, and H4K16ac (Multiple testing corrected p value shown on Fig7C). Differential histone modification was verified via western blot of independent MSCs and CA-MSCs (3 MSCs and 5 CAMSCs not used in the mass spec screen). All histone differences were confirmed with the exception of H3K79me1 (SupFigS5D,E). H3K27ac, H1K26me1 and H4K16ac are generally associated with transcriptional activation and were decreased in CA-MSCs. In contrast, H3K27me3 and H3K27me2 are associated with transcriptional repression and were increased in CA-MSCs. Three of the top six differential histone modifications are directly related to the PRC2: H3K27me2, H3K27me3 and H3K27ac. This supports a role for EZH2 differential targeting in CA-MSC formation.

Further supporting a role for differential EZH2 targeting in CA-MSCs, during cancer stimulated reprogramming of CA-MSCs, *EZH2* RNA expression is increased (Fig7D). This induction occurs at early stages of reprogramming with paracrine tumor signaling during indirect tumor cell: MSC co-culture (Fig7D), conditions which we demonstrated above induced DNA methylation changes in three of the five CA-MSC DMRs assessed in Figure 2. This induction is also seen with *in vivo* cancer stimulation in a xenograft mouse model as previous described (Coffman *et al*. 2019). This implies a role for EZH2 during the initial stages of CA-MSC formation.

### EZH2 inhibition in combination with *WT1* KD blocks the formation of a CA-MSC

Given our findings indicating altered EZH2 genomic targeting and known interaction between WT1 and EZH2/PRC2 targeting (Xu *et al*. 2011), we tested if EZH2 and WT1 directly interact in CA-MSCs. Using co-immunoprecipitation with an anti-EZH2 antibody in CA-MSCs, we demonstrated WT1 is pulled down with EZH2 (Fig7E). We then tested if EZH2 inhibition alone or in combination with WT1 KD would further decrease the development of a CA-MSC. To verify lack of toxicity, we performed viability studies with increasing dosage of an EZH2 inhibitor, GSK126. At doses of 20nM (Ki0.59nM, IC50 9.9 nM), there was no significant decrease in cell viability of tumor cells or MSCs (SupFig5A). Next, *WT1* KD and scrambled control MSCs were treated with 20nM GSK126 during cancer stimulation. GSK126 treatment of scrambled control MSCs lead to a 38% decrease in CA-MSC conversion. However, GSK126 treatment of *WT1* KD MSCs resulted in a 55% decrease in the conversion of *WT1* KD MSCs into CA-MSCs (Fig7F). Similar results were seen using a different EZH2 inhibitor, Tazemetostat (SupFig S5B). Collectively, this demonstrated WT1 and EZH2 jointly mediate the formation of a CA-MSC.

### EZH2 inhibition limits ovarian cancer metastasis

As EZH2 inhibition (EZH2i) disrupted the *in vitro* formation of CA-MSCs and CA-MSCs significantly enhanced metastasis, we tested if EZH2i impacted ovarian cancer metastasis. We used a previously described tail vein injection model of ovarian cancer which yields robust intra-abdominal metastasis (Coffman *et al*. 2016a). Mice were treated with GSK126 or vehicle control for one week prior to tail vein injection of OVCAR3 tumor cells and continued for two weeks after injection. Mice with vehicle control initial treatment were given GSK126 starting two weeks post-injection. EZH2i does not impact already established CA-MSCs (classifier score remains >0.97) (supFig S5C). Therefore, pretreatment/early treatment vs delayed treatment of GSK126 permitted comparison of tumor-mediated stromal reprogramming with and without EZH2i but controlled for the potential impact of EZH2i on tumor cell growth within an established TME. Four weeks post injection, all mice were sacrificed and necropsy was performed to identify areas of gross metastatic colonization. Strikingly, EZH2i decreased the rate of ovary malignant colonization by 60% (95% colonization in control group vs 35% colonization in EZH2i treated group) (Fig7G). Additionally, EZH2i decreased the amount of abdominal metastasis including bowel and mesentery involvement and ascites formation by 90%, 70% and 50% respectively. Rates of liver metastasis were decreased by 50% in the EZH2i group. Interestingly, rates of lung metastasis were equivalent (90% vs 100% in control vs EZH2i group). Additionally, lung tumor burden as measured by quantitative human specific PCR in whole lung tissue demonstrated equivalent tumor burden arguing against a direct cytotoxic effect of EZH2i which is consistent with the lack of *in vitro* toxicity (Fig7H).

## Discussion

Since described about a decade ago (Barrallo-Gimeno and Nieto 2005; Klymkowsky and Savagner 2009; Polyak and Weinberg 2009; Thiery *et al*. 2009; Yilmaz and Christofori 2009), partial transition to a mesenchymal phenotype in the epithelial compartment (EMT) in carcinomas has been well established as a hallmark of cancer with increased metastatic potential (Hanahan and Weinberg 2011). Here, we describe alterations in the other direction – mesenchymal to epithelial transition (MET) - in stromal progenitors of the TME in close contact with the carcinoma, that also contribute to cancer invasion and metastasis.

As stromal progenitor cells, CA-MSCs have a unique capacity to influence the formation of the TME. We previously demonstrated that CA-MSCs arise from cancer-mediated reprogramming of normal tissue MSCs creating a strongly pro-tumorigenic phenotype. Here we demonstrate that CA-MSCs have a unique epigenetic landscape different from both tumor cells and normal MSCs, characterized by DNA hypermethylation, altered chromatin accessibility and gain of repressive histone marks. The acquisition of DNA methylation at top DMRs during cancer stimulated reprogramming of MSCs into CA-MSCs validates an altered epigenetic landscape as a mitotically stable registrant of cancer stimulation. The majority of such differential DNA methylation occurs in enhancer regions. It has been previously shown that enhancers are the most dynamically utilized compartment (Calo and Wysocka 2013; Shlyueva *et al*. 2014; Sur and Taipale 2016) and closely related to cellular identity. Interestingly, at loci where CA-MSCs and MSC differed, the DNA methylation profile of CA-MSCs resembles benign FTE and sorted carcinoma while normal MSCs more closely resemble normal fibroblasts. Pathway analysis of differentially regulated genes demonstrated cell:cell adhesion pathways are upregulated in CA-MSCs. This is consistent with a more epithelial phenotype. While remaining mesenchymal in lineage, a partial MET appears to occur during CA-MSC formation. *In vitro* adhesion assays demonstrated CA-MSCs have a 3-fold increased capacity to bind tumor cells compared to normal MSCs also consistent with a MET phenotype. Importantly, this increase in tumor cell binding is critical for the pro-tumorigenic functions of CA-MSCs. CA-MSCs strikingly enhance ovarian cancer metastasis through a process of co-migration driven by direct CA-MSC:tumor cell contact. Indeed, CA-MSCs are abundant in ovarian cancer patient ascites as part of heterocellular clusters with tumor cells. This process of cancer cell and stromal cell co-migration is akin to the cancer “seed” traveling with its “soil” to enhance survival during metastasis and aid in distant colonization where the CA-MSC, as a stromal progenitor cell, may play a critical role in establishing the metastatic TME through stromal lineage differentiation.

Indeed, MET in CA-MSCs may be associated with increased differentiation capacity. De-differentiation of mesenchymal cells such as fibroblasts into induced pluripotent stem cells (iPSCs) requires MET (Li *et al*. 2010; Shu and Pei 2014). During embryogenesis, stem cells often transition from EMT to MET stages, and MET in stem cell reprogramming is associated with epigenetic changes, consistent with our findings (Wu *et al*. 2016). We previously reported that CA-MSCs compared to normal MSCs have enhanced “stemness” with increased colony formation, increased ALDH expression and greater differentiation potential consistent with greater pluripotency(McLean *et al*. 2011).

*WT1*, one of the most significantly upregulated genes in CA-MSCs, is an important mediator of MET during development and thought to be important in maintaining the potential to transition between EMT and MET (Miller-Hodges and Hohenstein 2012). Interestingly, WT1 has been shown to impact epigenetic regulation through directly binding the polycomb protein EZH2 and the DNA methyltransferases DNMT1 and DNMT3A (Xu *et al*. 2011; Szemes *et al*. 2013). Given the differential histone modifications (particularly related to PRC2 targets including H3K27me3) and DNA methylation changes in CA-MSCs, WT1 may be an important mediator of these epigenetic modifications. Indeed, co-IP confirmed WT1/EZH2 interaction in CA-MSCs, and knock down of WT1 in combination with EZH2 inhibition during cancer stimulation blocked the formation of a CA-MSC. Pharmacologic inhibition of EZH2, while not impacting tumor cell growth within the lung, significantly decreased the ability of ovarian cancer cells to form intra-abdominal metastasis. As metastasis is dependent on the colonization of a receptive TME, this supports the conclusion that EZH2 is important in the cancer stimulated epigenetic reprogramming of normal intra-abdominal MSCs to CA-MSCs.

Further, we demonstrated that the CA-MSC DNA methylation pattern persists throughout CA-MSC differentiation into other stromal elements including fibroblasts and adipocytes yet is not acquired during normal MSC differentiation. This points to the potential to develop a CA-MSC epigenetic signature which may be amplified throughout the stroma due to CA-MSC differentiation akin to generating a “field effect”. However, there may be opportunities to alter this stromal “field effect”. Omental MSCs derived from patients with a pathologic complete response to neoadjuvant chemotherapy fit a normal MSC profile indicating that either the CA-MSC phenotype is reversible or normal MSCs replace CA-MSCs after tumor death. Alternatively, these MSCs may be resistant to developing a CA-MSC phenotype and may be important in driving the excellent response to chemotherapy given the previously reported role of CA-MSCs in enhancement of chemotherapy resistance (Coffman *et al*. 2016b). Work currently under review in collaboration with the Buckanovich group demonstrated that CA-MSCs derived from metformin treated ovarian cancer patients have a methylation profile resembling normal MSCs supporting the ability to alter the development of a CA-MSC phenotype. Thus, understanding the mechanistic steps leading to CA-MSC development and the ability to reverse, replace or create MSCs resistant to reprogramming is essential to effective therapeutic targeting of the TME and represents a novel and powerful approach to ovarian cancer treatment.

## Acknowledgements

We would like to thank Marie Adams and the Genomics Core at Van Andel Institute (VAI) for the DNA methylation microarray service, and the VAI High Performance Computing Cluster supported by Zack Ramjan. We would like to also thank Dr. R. Buckanovich for providing cell lines. LGC is supported by NIH/NCI 7K08CA211362-02, Tina’s Wish Rising Star Grant, The Mary Kay Foundation Cancer Research Grant and DoD Ovarian Cancer, Omics Consortium Development Award. H. Shen is supported by NIH/NCI 1R37CA230748 and Ovarian Cancer Research Alliance’s Liz Tilberis Award (373933).

## Author contributions

Conceptualization, L.G.C and H.S.; Methodology, L.G.C, H.S, H.F, L.F, C.C, T.P, Y.W; Formal Analysis, L.G.C, H.S, H.F; Investigation L.G.C, L.F, C.C, H.F, H.A; Writing—Original Draft, L.G.C, H.S; Writing-Review & Editing, L.G.C, H.S, H.F, T.P, H.A; Funding Acquisition, L.G.C, H.S, I.S, T.W; Resources, L.G.C, H.S, T.P, I.S, T.W; Supervision, L.G.C, H.S, I.S, T.W

## Methods

### Tissue harvesting, culture

Patients samples were obtained in accordance with protocols approved by the University of Michigan’s IRB (HUM0009149) and University of Pittsburgh’s IRB (PRO17080326). Tissue was processed for DNA, RNA and protein isolation as previously described(Coffman *et al*. 2016b; Coffman *et al*. 2019). MSCs were isolated as previously described (McLean *et al*. 2011). Briefly, CA-MSCs were derived from surgical resection of human ovarian cancer involving the FT, ovary and/or omental metastatic deposits. Normal FT and omental MSCs were derived from surgical samples of women undergoing surgery for benign indications or risk reduction surgery. Cells were plated in supplemented MEBM, MSCs were selected for plastic adherence and cell surface marker expression (CD105, CD90, CD73 positive; CD45, CD34, CD14, CD19 negative). Adipocyte, osteocyte, and chondrocyte differentiation capacity was verified (following guidelines presented by the ISCT on the minimal criteria for defining multipotent mesenchymal stem cells (Dominici *et al*. 2006)) (SupFig6). To ensure lack of fibroblast contamination, cells were screened for fibroblast markers fibroblast surface protein (FSP) via flow cytometry and alpha smooth muscle actin via immunohistochemistry (SupFig6). Any cell line with a population >5% differing from above criteria were FACs purified. Fibroblast differentiation was performed as previously described by growing MSCs with connective tissue growth factor (100 ng/ml) and ascorbic acid (50 μg/ml) for 2 weeks (Coffman *et al*. 2019). MSCs were maintained in culture as previously described and used at passage 5 or below (McLean *et al*. 2011). All cell lines were tested and verified negative for mycoplasma (last test 9/2019).

### Cell Culture

Ovarian cancer cell lines CAOV3 and OVCAR3 were purchased through ATCC. OVSAHO and the primary patient cell line, PT412, were kind gifts from Dr. R. Buckanovich. All cells were grown in DMEM, 10% serum, 1% pen/strep. All cell lines were tested and verified negative for mycoplasma (last test 9/2019).

### DNA methylation array

DNA was isolated from CA-MSCs derived from omental metastasis in 6 patients with high grade serous ovarian cancer (HGSOC) and 1 patient with ovarian carcinosarcoma; 1 patient with high grade serous carcinoma involving the FT, 3 patients with high grade serous carcinoma involving the ovary. Samples from 4 patients with omental with initial radiographic involvement of HGSOC treated with neoadjuvant chemotherapy and pathologic complete response in the omentum, 5 benign patient FT samples, 1 set of bilateral FTs from a BRCA2 germline patient, 4 benign patient normal omental MSCs and 2 adipose MSCs purchased through ATCC. All normal MSCs were derived from patients undergoing surgery for benign indications. “A” and “B” indicate samples from the same patient from 2 distinct anatomic locations. A list of all samples with anatomic and histologic information is provided in supplemental table 1. All MSCs used for DNA methylation analysis were at passage 5 or below. DNA underwent bisulphite conversion followed by analysis on the Illumina MethylationEPIC Beadchip microarray through the Genomics core at the University of Michigan and Van Andel Research Institute.

### Assay for Transposase Accessible Chromatin with sequencing (ATACseq)

Nuclei were isolated from 4 CA-MSCs derived from omental metastasis of patients with high grade serous ovarian cancer, 2 normal omental MSCs and 2 adipose MSCs purchased from ATCC. All MSCs used for ATACseq analysis were at passage 5 or below. ATACseq was performed through Active Motif.

### Histone mass spectrometry

CA-MSCs and MSCs used for ATACseq were also used for mass spectrometry to measure the relative abundance of 80 histone modifications through Active Motif. Histones were extracted from frozen cell pellets and digested with trypsin. Samples were analyzed on a triple quadrupole (QdQ) mass spectrometer directly coupled with an UltiMate 3000 Dionex nan-liquid chromatography system. Three separate mass spec runs for each sample. Each modified and unmodified form of the monitored amino acid residue was quantified as the percentage of the total pool of modifications with mean and standard deviation reported.

### Quantitative real-time PCR

RNA was isolated with the RNeasy Mini Kit (Qiagen, Hilden, Germany) and on-column DNase treatment (Qiagen, Hilden, Germany). RNA concentration was determined with NanoDrop ND-1000 Spectrophotometer. cDNA was synthesized with the SuperScript III First-Strand Synthesis System for RT-PCR (Invitrogen, Grand Island, NY) as previously described [21]. SYBR green-based RT-PCR was performed using the 7900HT Sequence Detection System (Applied Biosystems, Foster City, California) and respective primers. The comparative Ct method was used for data analysis with *GAPDH* as the comparator gene.

### Immunoblotting

Cell pellets were homogenized in RIPA buffer (Pierce, Rockford, IL) with complete protease inhibitor (Roche, Basel, Switzerland). Insoluble material was removed by centrifugation at 16,000g at 4oC for 15mins. Protein concentrations were determined using the Bradford Protein Assay Kit (Bio-Rad, Hercules,CA). Equal amounts of protein were separated on 4-12% NuPAGE SDS gel (Invitrogen, Grand Island, NY) and transferred onto a PVDF membrane. Anti-H3K27me3, anti-H3K27me2, anti-H4K16ac, anti-H3K27ac, anti-H3K79me1, anti-H3, anti-H4, anti-WT1 and anti-EZH2 (1:1000 dilution, Active Motif) and anti-B-actin (1:10,000 dilution, Sigma-Aldrich, St. Louis, MO) were used. Bands were visualized using the ECL Kit (Pierce, Rockford, IL).

### Quantitative Methylation Specific PCR (qMSP)

*Bisulfite Conversion and DNA Yield* Bisulfite conversion of extracted cellular or tissue DNA was performed using the DNA Methylation-Lightning™ Kit (Zymo Research) according to manufacturer’s instructions and eluted into 40 µl DNA Elution buffer (Zymo Research). Post-bisulfite treatment DNA yields were quantified by MethyLight assay for the methylation-independent bisulfite converted (BSC) consensus ALU sequence, as previously described (Weisenberger *et al*. 2005), using primer and probe sequences: forward primer, 5’-GGT TAG GTA TAG TGG TTT ATA TTT GTA ATT TTA GTA-3’; reverse primer, 5’-ATT AAC TAA ACT AAT CTT AAA CTC CTA ACC TCA-3’, spanning a 98-bp locus, and 100nM probe, 5’-\56-FAM\ CCT ACC TTA ACC TCC C-3’ with Minor Groove Binder (MGB; ThermoFisher Scientific). PCR was performed using 10X Master Mix to yield a final volume of 25 µl and final working concentrations of 16.6mM (NH_4_)_2_SO_4_, 67mM Tris pH 8.8, 6.7mM MgCl_2_ 10mM β-mercaptoethanol, 200µM of each deoxynucleotide triphosphate (dNTP) and 0.04 U/µl of Platinum Taq polymerase (ThermoFisher Scientific). Cycling conditions were 95°C for 5 minutes, followed by 50 cycles of (95°C for 5 seconds, 60°C for 30 seconds and 72°C for 30 seconds). Standards for quantification were created by serial dilution of pre-quantified BSC human male DNA (Promega). Amplification reactions were performed in duplicate using 96 well-plates using a CFX96 Touch Real-time PCR Detection System (Bio-Rad).

*Locus-specific methylation analysis* qMSP assays (Lo *et al*. 1999) were developed and analytically-validated for five select loci: *SASH1*, *c13orf45*, *ELN*, *GATA6*, and *MARVELD2*, for which primer sequences and assay specifications are detailed in Supplementary Table 1. A standard curve was created using serial dilutions of BSC Epitect unmethylated control DNA (Qiagen) mixed with BSC CpG-Methylated HeLa Genomic DNA (New England BioLabs) (Lo *et al*. 1999).

*Data Analysis* MSP results were analyzed using the CFX Manager v3.1 (Bio-Rad) using regression to obtain the mean quantification cycle (Cq) for each respective sample. Percent methylation was calculated for each sample using the BSC-ALU and BSC Control DNA standard curves, according to the formula % methylation = (# locus copies methylated in sample)/(# ALU-calculated genomic copies in sample).

### CA-MSC encapsulation

CA-MSCs were encapsulated into 3% alginate as previously described (Schmitt *et al*. 2015). Briefly, 3% w/v sodium alginate was dissolved in PBS, filtered for sterility with a 0.2um filter. Cells were mixed with the alginate solution and added dropwise into a 5mM solution of calcium chloride to allow for gelation. Alginate beads were washed 2x with PBS prior to use in assays. Cell viability over time was verified by transection of the alginate capsule after 2 weeks incubation in standard culture media (DMEM, 10% FBS, 1% pen/strep) and re-isolation and growth of previously encapsulated CA-MSCs. Also, CA-MSCs lentivirally transduced with a secreted luciferase construct were encapsulated in alginate and the media was sampled for secreted luciferase overtime using the Ready-to-glow Secreted luciferase reporter system (Takara).

### Sphere assays

CA-MSCs transduced with lentiviral mT construct were mixed with lentiviral labeled GFP tumor cells (CAOV3, OVSAHO, OVCAR3) (as previously described (Coffman *et al*. 2019)) in a 1:1 ratio in ultra-low attachment 6 well plates grown in serum free supplemented MEBM as previously described (Coffman *et al*. 2019). Alternatively, CA-MSCs were added to the top of a 0.4um transwell system and tumor cells seeded in the bottom to prevent direct contact. After 5 days, the number of GFP+ spheres were counted. Experiments were repeated independently 3 times per tumor cell type.

### Migration/Invasion assays

8um transwells were coated with growth factor reduced Matrigel and allowed to solidify. GFP-tumor cells (CAOV3, OVSAHO, OVCAR3) with or without mT-CA-MSCs were added to the top of the transwell. Supplemented DMEM was added to the bottom of the transwell and cell migration/invasion through the coated membrane was quantified after 24hrs via fluorescent microscopy. Alginate encapsulated CA-MSCs were added in place of control CA-MSCs to determine the impact of preventing CA-MSC:tumor cell direct interaction. Experiments were repeated independently 3 times per tumor cell type.

### Adhesion assay

5×10∧4 MSCs or CA-MSCs were cultured overnight in 12 well-plate to form a mono-layer. 5×10∧4 fluorescently labeled tumor cells (OVSAHO, OVCAR3 or pt412) were then added. After 30 minutes, cells were washed twice with PBS and the attached tumor cells were counted using a fluorescence microscope and quantified as the number of adherent cells per low power field (10x). Experiments were repeated independently 3 times per tumor cell type.

### Cancer stimulation, cancer cell:MSC co-culture

MSCs were grown in a 1:1 ratio with ovarian tumor cells allowing for direct interaction or with use of a transwell system(0.4um) to separate the two cell types to allow only indirect co-culture as previously described (Coffman *et al*. 2019). Cells were grown in 1:1 mixture of CA-MSC media (supplemented MEBM) and tumor cell media (supplemented DMEM) for 5 days under hypoxic conditions (1% O2). Cells were isolated via FACs separation with fluorochrome labeled MSCs or tumor cells or removed from their respective transwell chamber.

### CA-MSC classifier

As previously described (Coffman *et al*. 2019), we created a regression model based on the expression of 6 genes which accurately distinguishes normal MSCs from CA-MSCs: Annexin A8-like protein 2 (ANXA8L2), Collagen Type XV Alpha 1 Chain (COL15A1), Cytokine Receptor Like Factor 1 (CRLF1), GATA Binding Protein 4 (GATA4), Iroquois Homeobox 2 (IRX2), and TGF-β2. Expression values are the delta CT values compared to GAPDH levels. The regression equation is: 1/(1+LOGIT((((B1*ANAX8L)+(B2*COL15)+(B3*CRLF1)+(B4*GATA4)+(B5*IRX2)+(B6*TGFb))+-7.62691)))); B1 = −0.00622, B2 = 0.175026, B3 = 0.886027, B4 = −0.34594, B5 = 0.416952, B6 = −0.00824

### Flow cytometric analysis

Cellular complexes were isolated from ovarian cancer patients’ ascites through centrifugation and serial washing steps with 2% FBS in PBS. The isolated complexes were enzymatically dissociated into a single cell suspension. Cells were then stained with anti-CD90, anti-CD73, anti-CD105 (Stem-cell Technology), and anti-EpCAM (Novus) antibodies for 20 minutes at RT. The percentage of CAMSC (CD90, 73, and 105+) and tumor cells (EpCAM+) were calculated on live cells gating.

### Immunofluorescence staining

Cellular complexes were isolated from ovarian cancer patients’ ascites as above and immobilized on slides using the cytospin centrifuge as previously described (Koh 2013). Then, the cellular complexes were fixed using methanol for 10 minutes. Immunofluorescence conjugated antibodies (anti-CD90, anti-CD73, anti-CD105 (Stem-cell Technology), and anti-EpCAM (Novus)) were added after permeabilize the cellular complexes with 0.1% Triton 100X. The stained cellular complexes were examined using Nikon A1 confocal microscope.

### Orthotopic ovarian cancer mouse model

5×10∧5 OVCAR3 cells were injected alone (n=5) or with control CA-MSCs (5×10∧5 cells) (n=5) or encapsulated CA-MSCs (5×10∧5 cells) (n=4) into the ovarian bursa of NSG mice. Mouse weight and health were monitored over time and mice were sacrificed when the first group of mice met endpoint criteria of >10% weight loss and necropsy was performed.

### Tail vein injection model

5×10∧5 OVCAR3 cells were injected into NSG mice (n=10 per group). Mice in the EZH2 inhibition group were treated with 300mg/kg of GSK126 (selleckchem) 3 x weekly for 1 week prior to injection and then 150mg/kg 3 x weekly for 2 weeks immediately after injection. Mice in the control group underwent parallel treatment with vehicle control and were treated with 150mg/kg 3x weekly for 2 weeks starting 2 weeks after treatment. Mice were euthanized at 4 weeks post tumor cell injection and necropsy performed. Quantitative human specific PCR was performed on whole lung isolates as previously described (Alcoser *et al*. 2011).

### Transcriptome data re-processing and analysis

Raw mRNA sequencing data (FASTQ format) of 4 normal omental MSCs and 10 ovarian CA-MSCs, using Illumina TruSeq RNA Sample Preparation V2 kit (Illumina San Diego, CA, USA) and Illumina HiSeq2000 instrument 100 bp PE sequencing, were download through NCBI’s Gene Expression Omnibus (GEO) with GEO Series accession number GSE118624 (https://www.ncbi.nlm.nih.gov/geo/query/acc.cgi?acc=GSE118624) (Coffman *et al*. 2019). Each raw sequencing file was aligned to the human reference genome (GRCh37) using STAR version 2.7 (Dobin *et al*. 2013), with default settings. Estimation of gene-level abundance was carried out using RSEM version 1.3.1 (Li and Dewey 2011). Raw reads count from RSEM were further normalized using R package *edgeR* (function cpm), and log-transformed for downstream analysis, with one CA-MSC sample (GSM3335697) removed due to potential tumor cell contamination.

Differential gene expression analysis was carried out using R package Limma (Ritchie *et al*. 2015). Specifically, differentially expressed genes were identified based on a p-value less than 0.05, and an absolute log-transformed fold change greater than 1.5. Followed by functional enrichment analysis using R package clusterProfiler (Yu *et al*. 2012), with FDR cutoff at 0.05 for KEGG pathway, Gene Ontology (Biological Process), and Gene Set Enrichment Analysis (GSEA). In particular for GSEA, genes were ranked based on log-transformed fold changes by comparing sample group of CA-MSC to MSC.

### DNA methylation data processing and QC

Raw IDATs files were processed using R package SeSAMe (Zhou *et al*. 2018) with noob background correction, non-linear dye bias correction, and non-detection masking (any data point not significantly different from background was replaced with NA). Probes with design issues are also masked (Zhou *et al*. 2017). DNA methylation beta values, ranging from 0 to 1 (with “0” indicating fully unmethylated and “1” fully methylated), were calculated as quantitative percentage of methylated signals over both methylated and unmethylated signals. All IDATs and processed beta values have been uploaded and made available through Gene Expression Omnibus (GEO) under the accession number GSE138072.

SNP probes (‘rs’ probes) were used to examine potential sample swaps which can occur in genomic studies. No such swap was identified. DNA Methylation beta values for three MIR141/200C promoter probes (“cg12161331”, “cg18185189”, “cg19794481”) were examined to track mesenchymal content within each sample. MIR141/200C is master regulator for epithelial/mesenchymal phenotype transition, and this process is controlled by its promoter methylation state (Vrba *et al*. 2010). Methylation level at these three probes used highly correlated with mesenchymal content in flow sorting results (George *et al*. 2018). Two samples had a failure rate of >40% with our stringent background masking (Zhou *et al*. 2018) likely due to DNA degradation, but were kept in the study as the remaining data points were passed the very stringent filter and represented high quality data.

### Unsupervised and supervised DNA Methylation analysis

Unsupervised hierarchical clustering was performed on top variable CpG probes (N=10,000, filtered by standard deviations) across all CA-MSC and MSC samples measured on the EPIC array with the R function *hclust*(). Uniform Manifold Approximation and Projection (UMAP) was performed with the R package *uwot.*.

For global comparison of CAMSC/FTE/MSCs, different probe categories are defined as such: 1) CA-MSC and FTE shared hyper-methylated probes: mean methylation difference > 0.4 in both CA-MSC to MSC, and FTE to MSC comparisons; 2) CA-MSC-specific hyper-methylated probes: mean methylation difference > 0.4 when comparing CA-MSC to MSC, but absolute mean methylation difference < 0.1 when comparing FTE to MSC; 3) FTE-specific hyper-methylated probes: mean methylation difference > 0.4 when comparing FTE to MSC, but absolute methylation difference < 0.1 when comparing CA-MSC to MSC; 4) CA-MSC and FTE shared hypo-methylated probes: mean methylation difference < −0.4 in both CA-MSC to MSC, and FTE to MSC comparisons; 5) CA-MSC-specific hyo-methylated probes: mean methylation difference < −0.4 when comparing CA-MSC to MSC, but absolute methylation difference < 0.1 when comparing FTE to MSC; 6) FTE-specific hypo-methylated probes: mean methylation difference < −0.4 when comparing FTE to MSC, and absolute mean methylation difference < 0.1 when comparing CA-MSC to MSC.

Differentially methylated cytosines (DMCs) were calculated using R package DMRcate (Peters *et al*. 2015), by comparing sample group of CA-MSC to MSC, based on its default FDR cutoff of 0.5. DMRcate does not tolerate missing values (NA), and therefore we removed probes with any NA in any of the samples, leaving a total of ∼401,000 probes for any analyses involving DMR/DMC calling. DMCs were split into hypermethylated (hyper-) and hypomethylated (hypo-) DMCs, by comparing group of CA-MSC to MSC. Enrichement or depletion of various genomic features (annotated with R package *annotatr* for each probe) in hyper- and hypo DMCs were measured as odds ratios, together with significance level calculated using Chi-square test. DMCs were mapped to genes captured by mRNA sequencing, if they were located within gene promoter regions defined as +/-1.5 kb away from transcription start sites (TSSs). Annotation of CpG probes against transcription factor binding sites (TFBSs) were adopted from our previous study (Zhou *et al*. 2017). We then performed enrichment analysis of TFBSs over differentially methylated probes at distal regulatory elements (i.e. potential enhancers). Distal probes were defined as probes located within TFBSs, but not overlapping with +/-2 kb flanking regions surrounding TSSs. For each set of hyper- or hypo-DMCs, hypergeometric test was applied to calculate the enrichment of binding sites for each TF within the set of DMCs within all distal probes. Significance cutoff was made at p=1e-3 after false discovery rate (FDR) correction.

### ATACseq data processing and analysis

Quality control was carried out using the PEPATAC pipeline adopted by the TCGA ATAC-seq project (Corces *et al*. 2018). We followed current ENCODE QC standards and excluded four samples due to insufficient transcription start site (TSS) enrichment value (less than 6), therefore were removed from further analysis. Next, raw sequencing reads were mapped to the human reference genome (GRCh37) using Burrows-Wheeler Alignment tool (BWA) with default settings (Li and Durbin 2009), followed by peak calling with software MACS2 (Zhang *et al*. 2008). Specifically, parameters of --nomodel and -f BAMPE were specified in calling open chromatin regions, for pileup of the whole fragments. Sample-based open peaks were then merged across all the samples, and fed into R package DESeq2 (Love *et al*. 2014) to generate a normalized reads count matrix against effective library sizes. Log-transformed reads count were used for further differential peak analysis, with reads counts of replicates from the same sample averaged. Principal component analysis (PCA) was performed, based on the normalized reads count matrix. Peak summit centered heatmap per sample was generated with deepTools using normalized 10 base pair bin-based ATACseq signal (Ramírez *et al*. 2016).

### Public available data resources

DNA methylation data (Illumina HM450 array) of ovarian stroma samples (N=4; samples annotated as ovary and pathologist evaluation of available slides confirms ovarian stroma histology) were downloaded from the Genomic Data Commons (GDC) Data Portal (https://portal.gdc.cancer.gov), together with that of Uterine Carcinosarcoma (UCS; N=57) and Sarcoma (SARC; N=261). Pure, uncultured fallopian tube epithelium samples (FTE; N=4) were downloaded through GEO database, with one sample GSM2146822 taken from Series GSE81224 (Klinkebiel *et al*. 2016), and the other three samples (GSM1606969, GSM1606970, and GSM1606973) from Series GSE65820 (Patch *et al*. 2015). Other FTE samples in the same studies either had insufficient epithelial purity or had undergone *in vitro* culturing and therefore not included. All DNA methylation microarray data were reprocessed from raw IDATs in the same way as described above for the MSCs. Gapped H3K27me3 chromatin peak files of MSC-derived from adipose, bone marrow, chondrocyte, and H1 stem cell were downloaded through the NIH Roadmap Epigenomics projects (http://www.roadmapepigenomics.org/data/). EZH2 target genes were then defined as genes overlapping with H3K27me3 marked regions shared by all four different cell types.

## Supplemental information title and legend

**Supp Figure S1.**
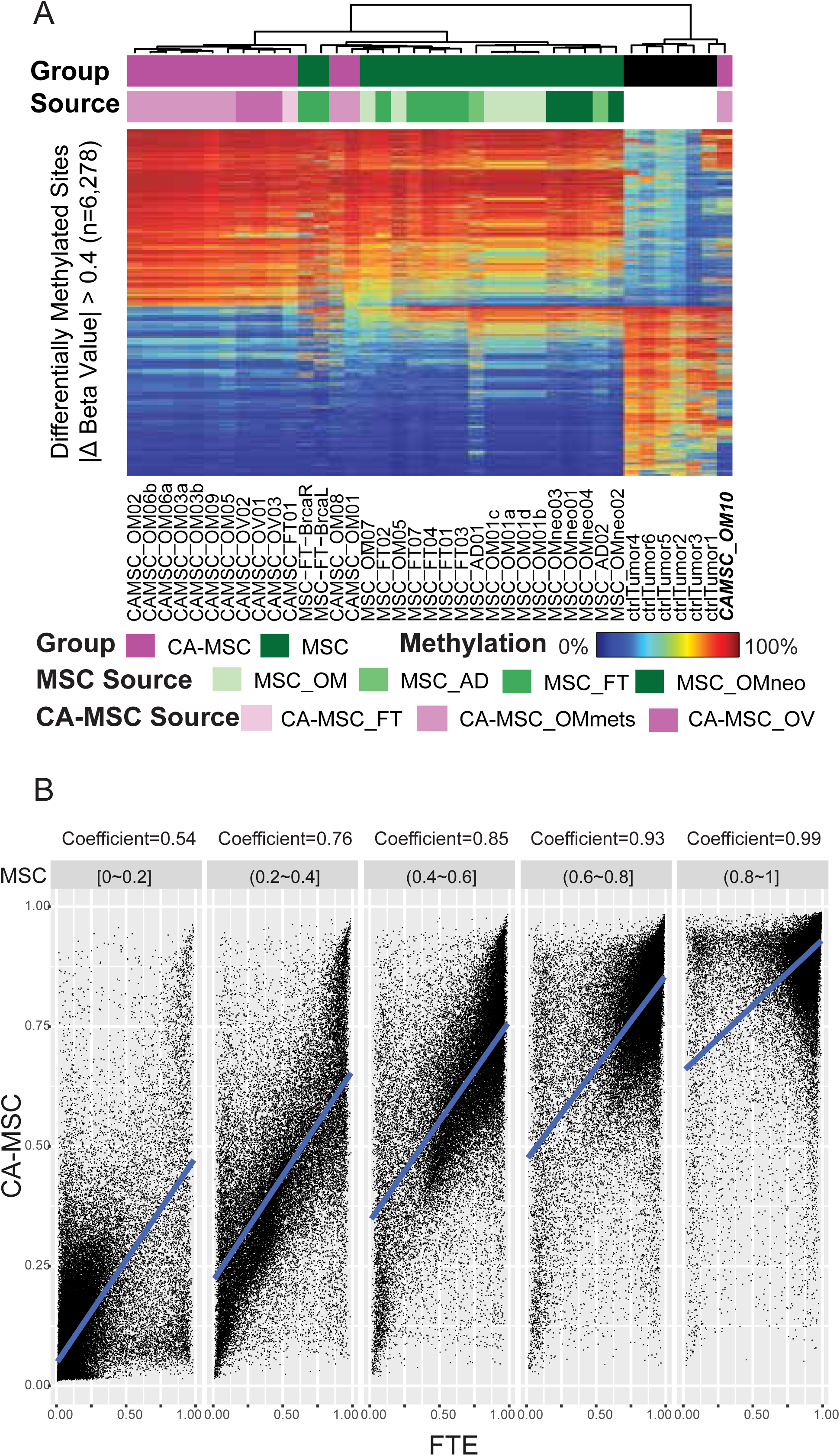
A. DNA methylation heatmap of top differentially methylated loci (FDR<0.05, and absolute group beta value difference > 0.4) between tumor samples and CA-MSCs. Probes are plotted as rows, with samples plotted as columns. Sample group of tumor, MSC, and CA-MSC is plotted as column annotation, together with sample isolation source if samples are MSCs or CA-MSCs. Sample CA-MSC_OM10 with heavy epithelial contamination is highlighted. B. Dot plot of probe-level similarity between CA-MSC and fallopian tube epithelium (FTE) samples, stratified by their methylation level in MSCs. Linear regression lines are in blue, with corresponding coefficient indicated on top of each dot plot. A blue-to-red gradient indicates a beta value of 0-1 (DNA methylation level of 0% to 100%).

**Supp Figure S2.**
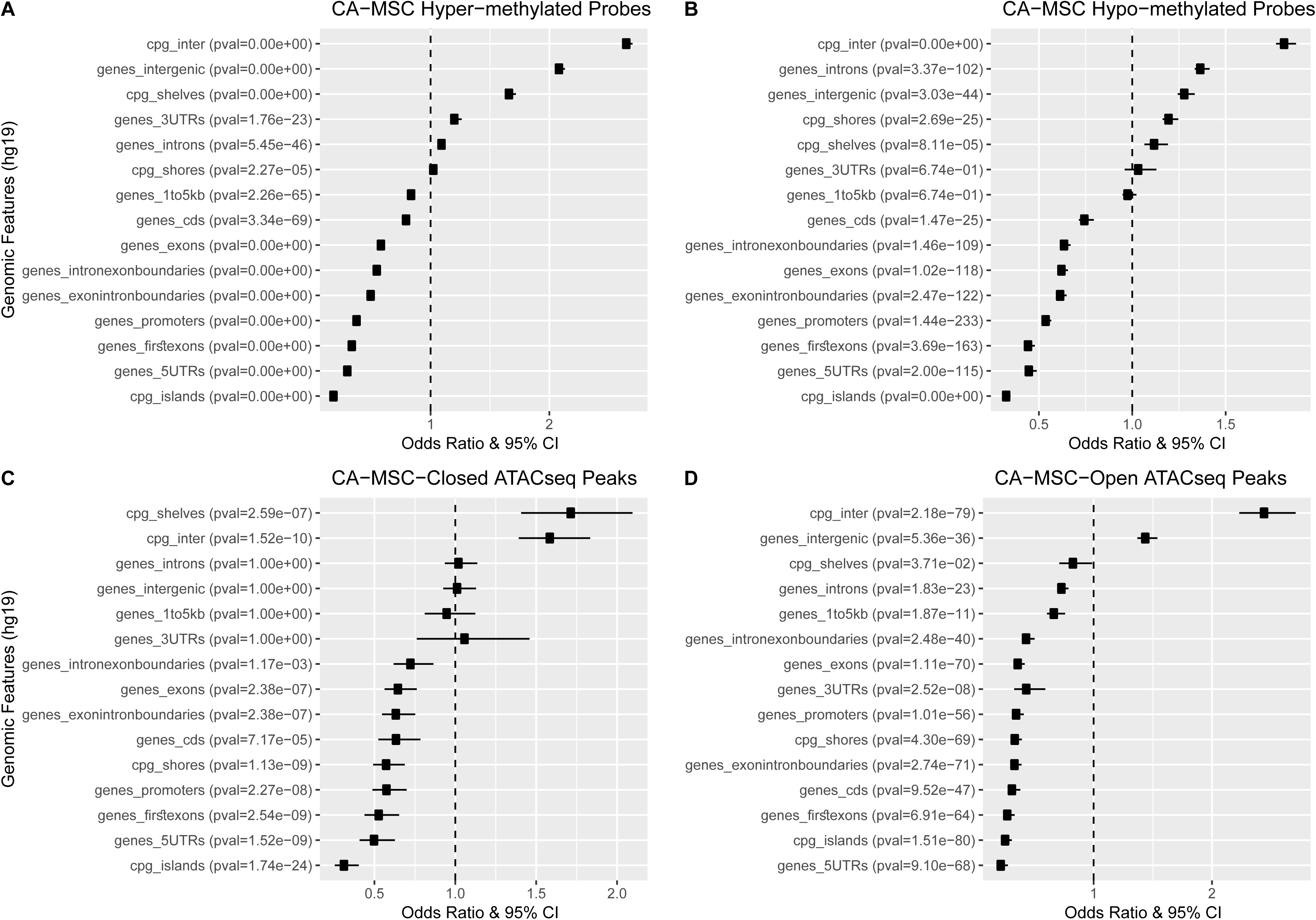
A. Enrichment of CA-MSC hyper-methylated loci against different genomic features of CpG and genic annotations. CpG annotations include open sea (cpg_inter), CpG shores (cpg_shores), CpG shelves (cpg_shelves), CpG islands (cpg_islands). Genic annotations contain 1-5Kb upstream of the TSS (genes_1to5kb), the promoter (< 1Kb upstream of the TSS; genes_promoters), 5’UTR (genes_5UTRs), first exons (genes_firstexons), exons (genes_exons), introns (genes_introns), CDS (genes_cds), 3’UTR (genes_3UTRs), and intergenic regions (the intergenic regions exclude the previous list of annotations; genes_intergenic).

**Supp Figure S3.**
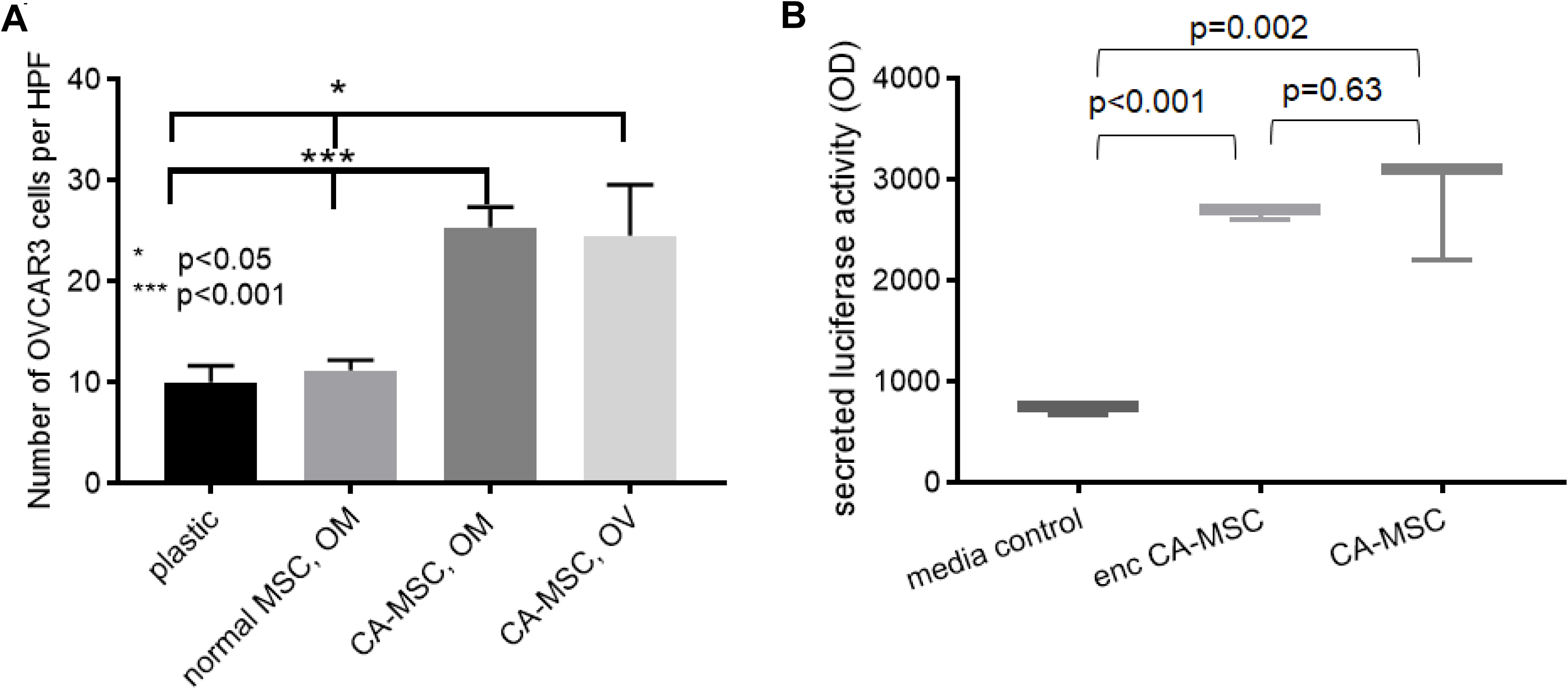
A. Adhesion of a third cell line in addition to what was shown in Figure 5, OVCAR3 cells to plastic, normal omentum (OM) MSCs, CA-MSCs derived from the OM and ovary (OV) demonstrates increased tumor cell adhesion to CA-MSCs. Results quantified by counting fluorescent tumor cells per high power field (HPF). Mean & SEM of 3 independent experiments is represented. B. Luciferase secretion of encapsulated (enc.) CA-MSCs. CA-MSCs transduced with a secreted luciferase construct were encapsulated into alginate and the amount of secreted luciferase after 5 days was compared. Alginate encapsulation did not alter CA-MSC protein secretion.

**Supp Figure S4.**
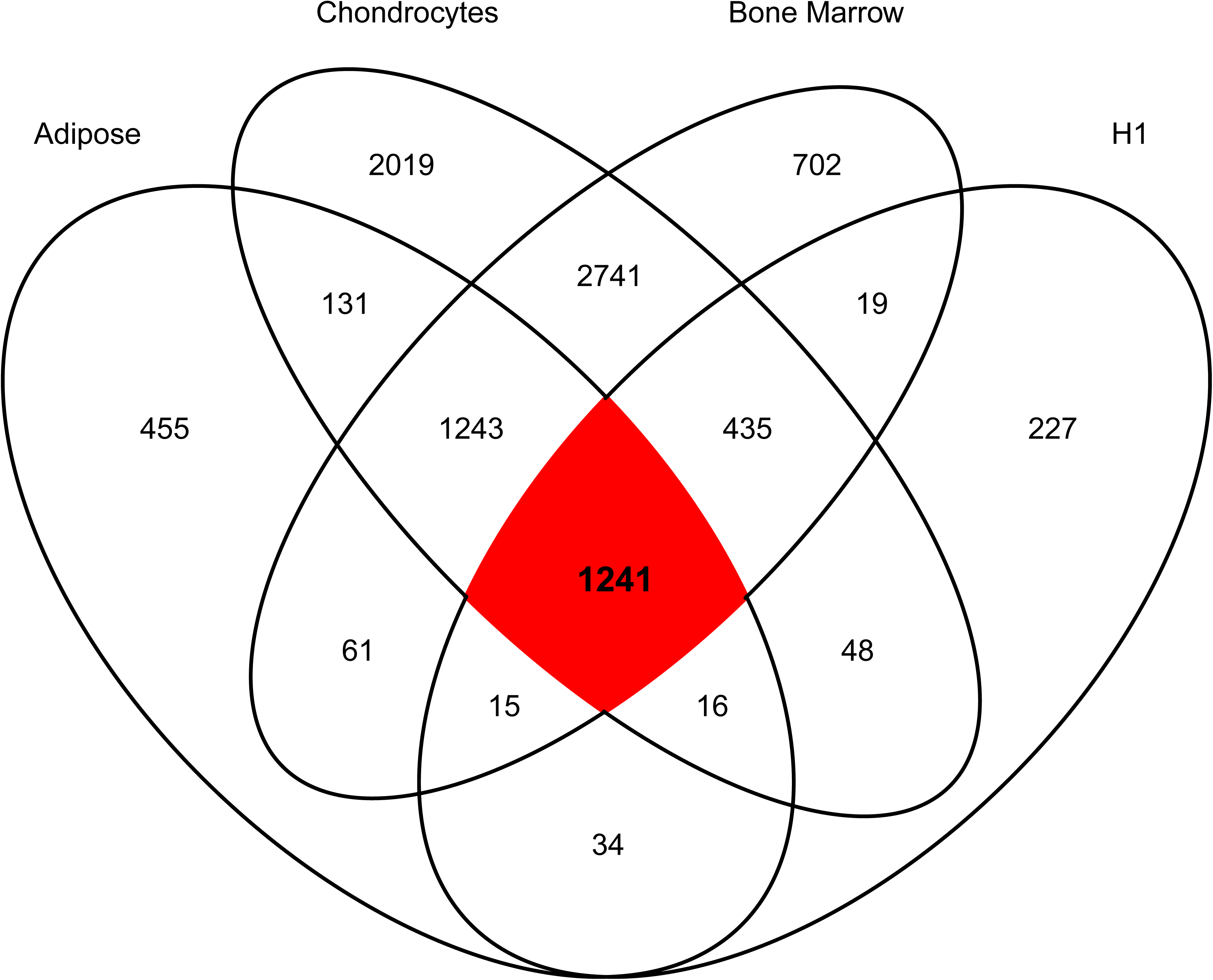
Venn diagram showing the H3K27me3 peak overlaps among MSCs derived from different sources of adipose, chondrocytes, bone marrow, and H1 cell line. The common set of 1241 peak regions are applied to map the EZH2 targets (as in red), if they are located within their transcriptional regions plus 2 kb upstream flanking regions of their TSSs.

**Supp Figure S5.**
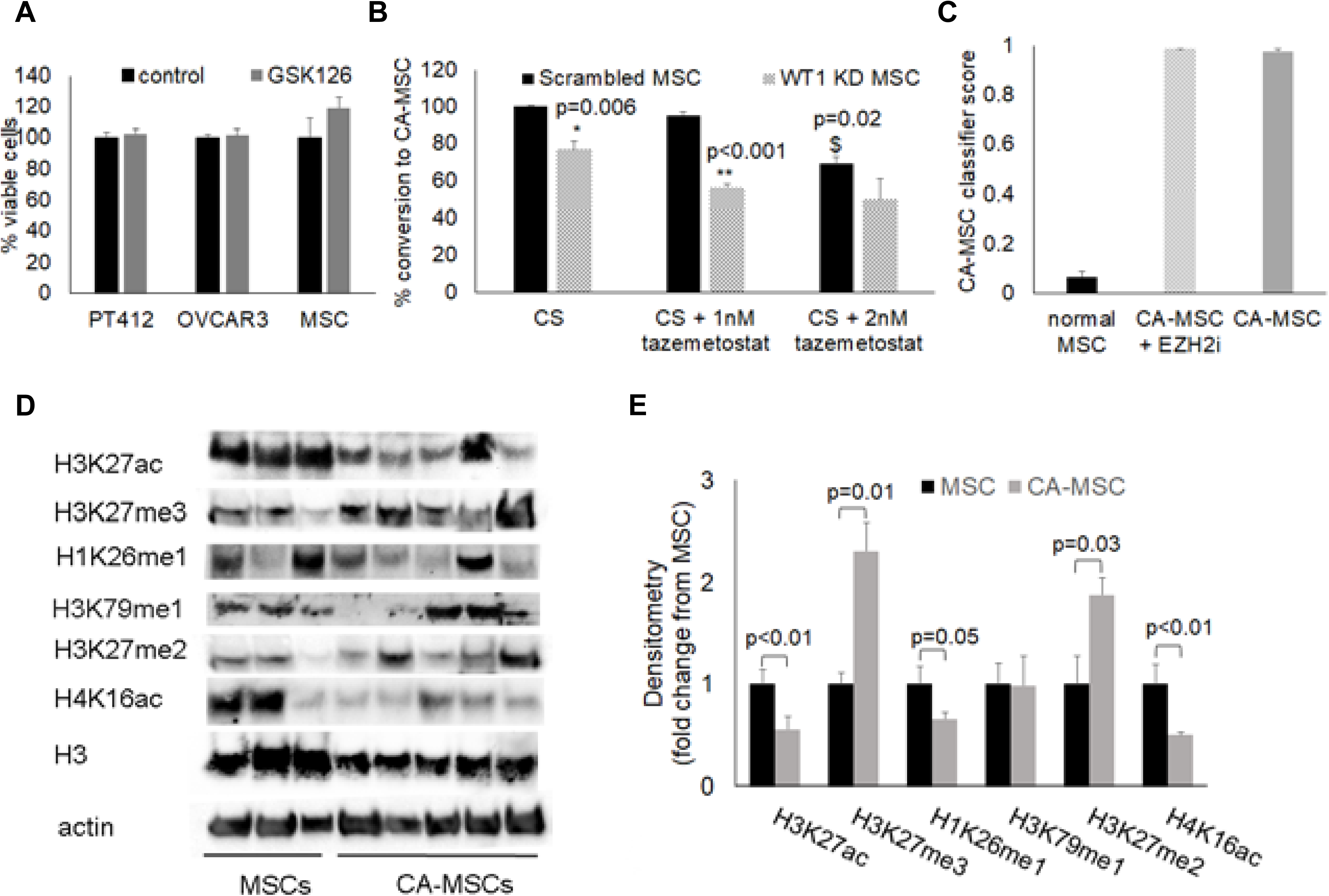
A. Viability of tumor cells PT412 and OVCAR3 and normal MSCs after 5 days treatment with 20nM GSK126 compared to vehicle control. B. Quantification of the percent of conversation to CA-MSC in WT1 KD MSCs or scrambled control after cancer stimulation (CS) with hypoxic direct cancer cell co-culture with and without a different EZH2 inhibitor, Tazemetostat. Values are normalized to scrambled control conversion based on CA-MSC classifier score. *, ** = comparison to treatment group scrambled control; $ = comparison to CS scrambled control without Tazemetostat C. Measurement of the CA-MSC classifier score in normal MSCs, CA-MSCs and CA-MSCs treated with EZH2i (GSK126 or Tazemetostat) demonstrating EZH2i does not impact the classifier score of already established CA-MSCs. For all in vitro experiments, mean & SEM of 3 independent experiments is represented. D. Western blot verification of differential histone modifications identified by mass spec. E. Densitometry quantification of histone modification western blots, compared to actin control.

**Supp Figure S6.**
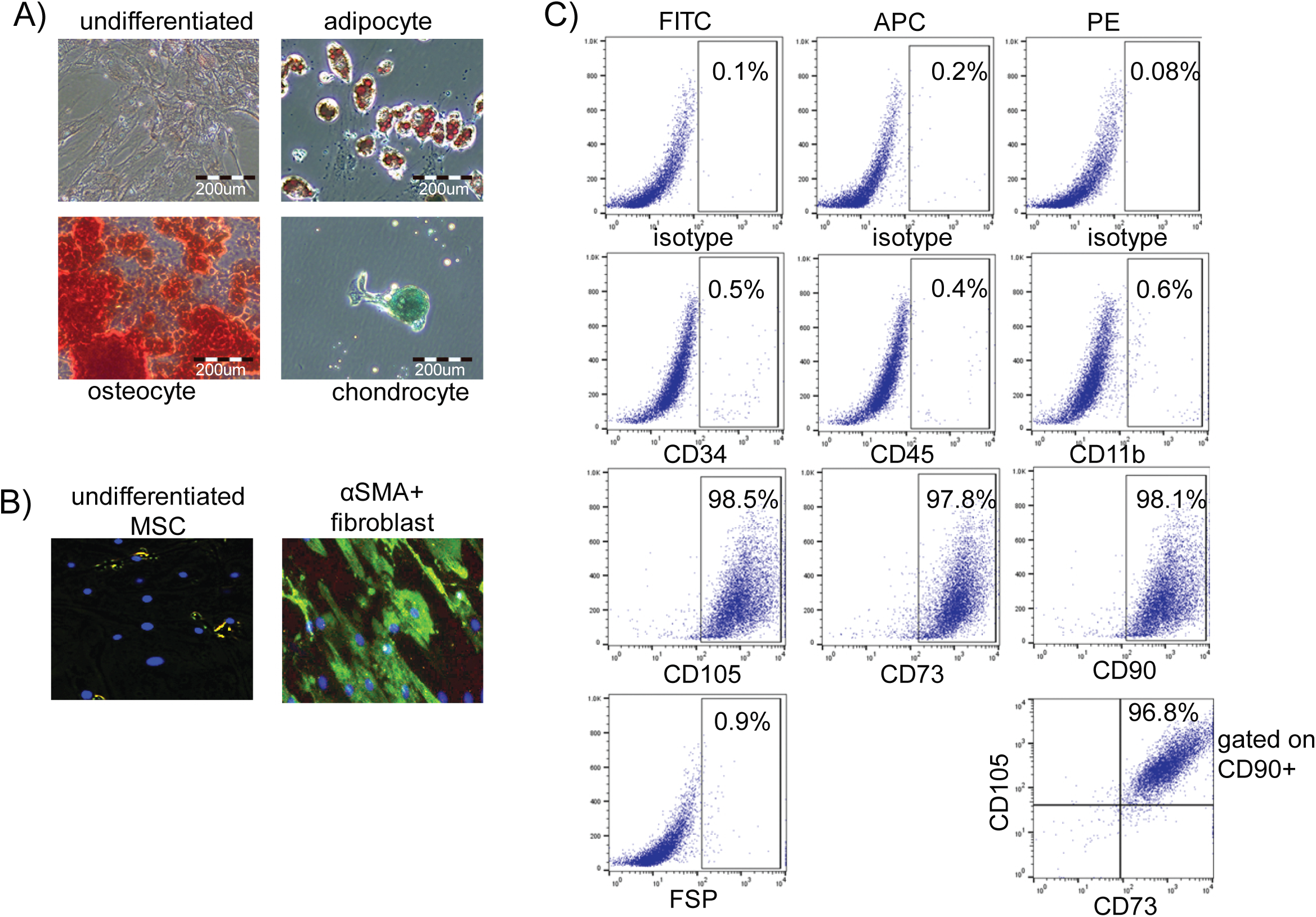
A. Immunohistochemistry of undifferentiated and differentiated MSCs (adipocyte: oil red o; osteocyte: alizarin red; chondrocyte: alcian blue). B. Immunofluorescence of alpha-smooth muscle actin (aSMA) in undifferentiated and fibroblast differentiated MSCs. C. Representative flow cytometry plots characterizing MSCs based on CD105,73,90 positivity and CD34,45,11b negativity and negativity for the fibroblast marker, fibroblast surface protein (FSP).

**Supp. Table 1.** Sample list summary

| SampleName | SampleID | PatientID | Sample Group | SampleEPIC | Source | SourceCode |
| --- | --- | --- | --- | --- | --- | --- |
| CAMSC_OM01 | 493_cams<br>c | s493 | CAMSC | 201557520003<br>_R03C01 | CA-MSCs derived from the omental metastasis of patients with high grade serous ovarian cancer | CAMSC-OMmets |
| CAMSC_OM02 | 549_cams<br>c | s549 | CAMSC | 201557520040<br>_R03C01 | CA-MSCs derived from the omental metastasis of patients with high grade serous ovarian cancer | CAMSC-OMmets |
| CAMSC_OM03<br>a | CAMSC1 | s0009 | CAMSC | 200516380137<br>_R01C01 | CA-MSCs derived from the omental metastasis of patients with high grade serous ovarian cancer | CAMSC-OMmets |
| CAMSC_OM03<br>b | CAMSC2 | s0009 | CAMSC | 200516380137<br>_R02C01 | CA-MSCs derived from the omental metastasis of patients with high grade serous ovarian cancer | CAMSC-OMmets |
| CAMSC_OM05 | CAMSC3 | s0003 | CAMSC | 200516380137<br>_R03C01 | CA-MSCs derived from the omental metastasis of patients with high grade serous ovarian cancer | CAMSC-OMmets |
| CAMSC_OM06<br>a | CAMSC6 | s0006 | CAMSC | 200516380137<br>_R06C01 | CA-MSCs derived from the omental metastasis of patients with high grade serous ovarian cancer | CAMSC-OMmets |
| CAMSC_OM06<br>b | CAMSC7 | s0006 | CAMSC | 200516380137<br>_R07C01 | CA-MSCs derived from the omental metastasis of patients with high grade serous ovarian cancer | CAMSC-OMmets |
| CAMSC_OM08 | CAMSC8 | s0008 | CAMSC | 200516380137<br>_R08C01 | CA-MSCs derived from the omental metastasis of patients with ovarian carcinosarcoma | CAMSC-OMmets |
| CAMSC_OM09 | CAMSC9 | s0009 | CAMSC | 200526210119<br>_R01C01 | CA-MSCs derived from the omental metastasis of patients with high grade serous ovarian cancer | CAMSC-OMmets |
| CAMSC_OM10 | 406_cams<br>c | s406 | CAMSC | 201557520040<br>_R02C01 | CA-MSCs derived from the omental metastasis of patients with high grade serous ovarian cancer | CAMSC-OMmets |
| CAMSC_FT01 | 19-175<br>FT.camsc | s19-175 | CAMSC | 203020780110<br>_R04C01 | CAMSCs derived from fallopian tube invovled with high grade serous ovarian cancer | CAMSC-FT |
| CAMSC_OV01 | 19-90<br>ov.camsc | s19-90 | CAMSC | 203020780110<br>R05C01 | CAMSCSs derived from ovary involved with high grade serous ovarian cancer | CAMSC-OV |
| CAMSC_OV02 | 19-95<br>ov.camsc | s19-95 | CAMSC | 203020780110<br>R06C01 | CAMSCSs derived from ovary involved with high grade serous ovarian cancer | CAMSC-OV |
| CAMSC_OV03 | 19-263<br>ovcamsc | s19-263 | CAMSC | 203038250154<br>R06C01 | CAMSCSs derived from ovary involved with ovarian cancer | CAMSC-OV |
| MSC_OMneo01 | 18-0030<br>neo n.om | s18-0030 | MSC | 203020780110<br>_R08C01 | MSCs derived from omentum that was initially involved with high grade serous ovarian cancer but had a complete pathologic response to neoadjuvant chemotherapy | MSC-OMneo |
| MSC_OMneo02 | 18-0054<br>neo n.om | s18-0054 | MSC | 203013220061<br>_R01C01 | MSCs derived from omentum that was initially involved with high grade serous ovarian cancer but had a complete pathologic response to neoadjuvant chemotherapy | MSC-OMneo |
| MSC_OMneo03 | 18-0048<br>neo n.om | s18-0048 | MSC | 203013220061<br>_R02C01 | MSCs derived from omentum that was initially involved with high grade serous ovarian cancer but had a complete pathologic response to neoadjuvant chemotherapy | MSC-OMneo |
| MSC_OMneo04 | 19-107<br>n.om<br>(neo) | s19-107 | MSC | 203013220061<br>_R03C01 | MSCs derived from omentum that was never involved with cancer (stage II ovarian cancer) but patient had neoadjuvant chemotherapy | MSC-OMneo |
| MSC_FT01 | 19-120<br>n.FT | s19-120 | MSC | 203013220061<br>_R04C01 | MSCs derived from normal fallopian tube | MSC-FT |
| MSC_FT02 | 19-197<br>n.FT | s19-197 | MSC | 203013220061<br>_R05C01 | MSCs derived from normal fallopian tube | MSC-FT |
| MSC_FT03 | 19-128<br>n.FT | s19-128 | MSC | 203013220061<br>_R06C01 | MSCs derived from normal fallopian tube but with hydrosalpinx | MSC-FT |
| MSC_FT04 | AL n.FT | s0007 | MSC | 203013220061<br>_R07C01 | MSCs derived from normal fallopian tube | MSC-FT |
| MSC-FT-BrcaL | 19-257<br>LFT<br>BRCA2 | s19-257 | MSC | 203038250154<br>_R02C01 | MSCs derived from left fallopian tube with BRCA2 mutation | MSC-FT |
| MSC-FT-BrcaR | 19-257<br>RFT<br>BRCA2 | s19-257 | MSC | 203038250154<br>_R03C01 | MSCs derived from right fallopian tube with<br>BRCA2 mutation | MSC-FT |
| MSC_FT07 | 19-223 #2<br>nFT | s19-223 | MSC | 203038250154<br>_R04C01 | MSCs derived from normal fallopian tube | MSC-FT |
| MSC_AD01 | amsc_112<br>5 | s1125 | MSC | 201557520040<br>_R04C01 | MSCs derived from adipose purchased<br>through ATCC | MSC-AD |
| MSC_AD02 | amsc_211<br>8 | s2118 | MSC | 201557520003<br>_R05C01 | MSCs derived from adipose purchased<br>through ATCC | MSC-AD |
| MSC_OM01a | MSC1 | s0001 | MSC | 200526210119<br>_R02C01 | MSCs derived from normal omental tissue<br>undergoing surgery for benign indications | MSC-OM |
| MSC_OM01b | MSC2 | s0001 | MSC | 200526210119<br>_R03C01 | MSCs derived from normal omental tissue<br>undergoing surgery for benign indications | MSC-OM |
| MSC_OM01c | MSC5 | s0001 | MSC | 200526210119<br>_R06C01 | MSCs derived from normal omental tissue<br>undergoing surgery for benign indications | MSC-OM |
| MSC_OM01d | MSC6 | s0001 | MSC | 200526210119<br>_R07C01 | MSCs derived from normal omental tissue<br>undergoing surgery for benign indications | MSC-OM |
| MSC_OM05 | 18-691<br>n.om CC | s18-691 | MSC | 203013220061<br>_R08C01 | MSCs derived from normal omentum from<br>patient with a stage II clear cell endometrial<br>cancer | MSC-OM |
| MSC_OM05b | 18-691<br>n.om CC | s18-691 | MSC | 203038250154<br>_R01C01 | MSCs derived from normal omentum from<br>patient with a stage II clear cell endometrial<br>cancer | MSC-OM |
| MSC_OM07 | 19-319<br>n.om | s19-319 | MSC | 203038250154<br>_R05C01 | MSCs derived from normal omentum | MSC-OM |

**Supp. Table 2.**
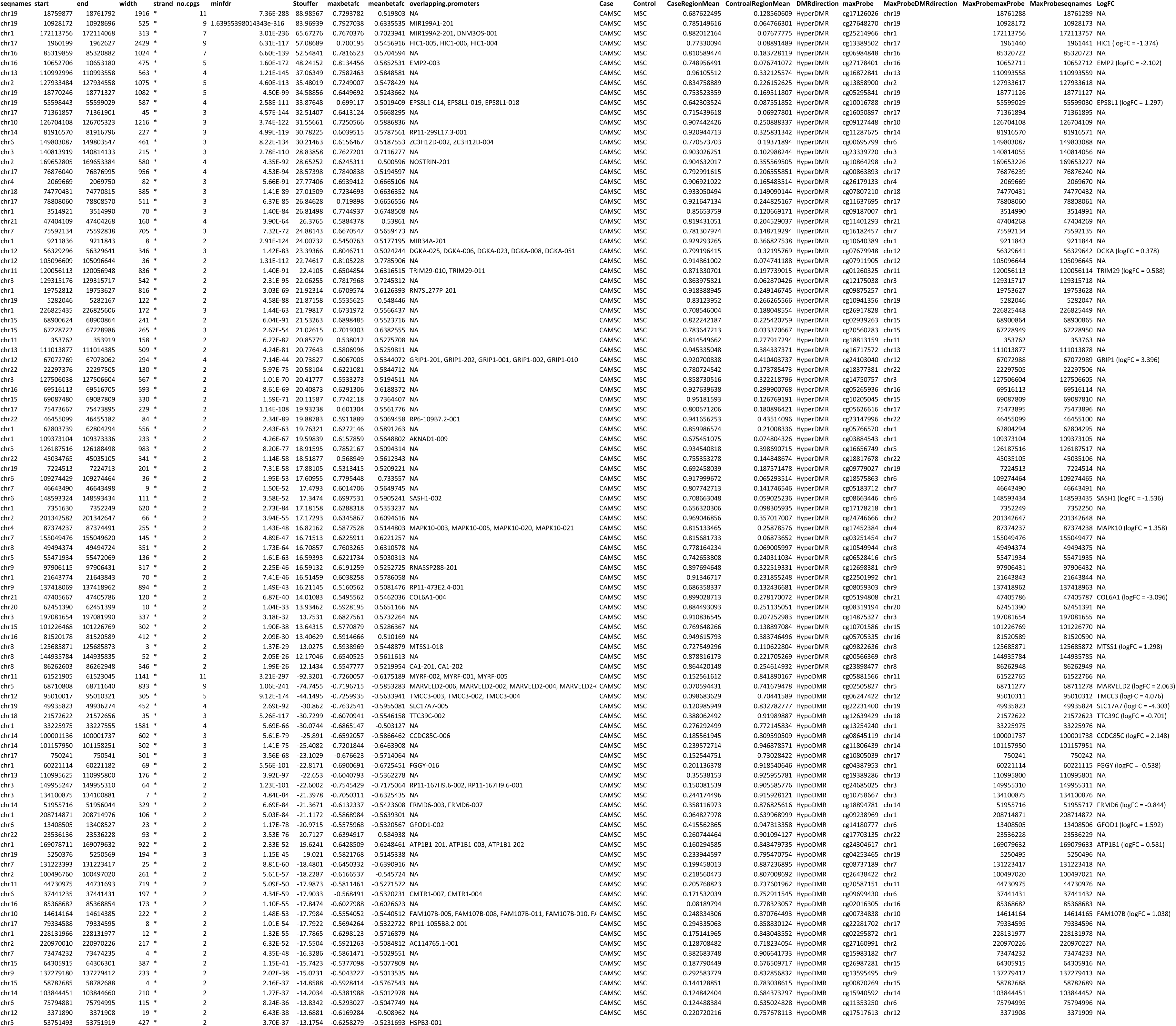
Most significant differentially methylated regions (DMRs) between CA-MSCs and normal MSCs (ranked P-value followed by mean beta value levels).

